# Conformational dynamics of the active state of β-arrestin 1

**DOI:** 10.1101/2025.06.10.658911

**Authors:** Van Ngo, Wesley B. Asher, Jonathan A. Javitch, Lei Shi

## Abstract

β-arrestins (βarr) regulate the signaling and trafficking of G protein-coupled receptors (GPCRs) in numerous physiological processes and have been implicated in various diseases. Structural and kinetic insights into how ligand-mediated GPCR activation drives βarr coupling and activation remain limited, with the binding mechanism of the phosphorylated GPCR C-terminal tails, such as that of the vasopressin receptor-2 (V2Rpp), and the conformation of the entire βarr tail in the active state still unknown. Here, we simulated both the basal and V2Rpp-bound states of βarr1 with temperature replica-exchange molecular dynamics (TREMD) simulations to probe the activation mechanism of βarr1. Compared to conventional MD, our TREMD simulations, employing an unprecedented 200-replica setup, significantly broadened conformational sampling while preserving the basal state. Our analysis showed that, without the bound Fab30 antibody fragment, the main body of V2Rpp-bound βarr1 tended to transition toward the basal conformation; however, binding of V2Rpp in the N-domain groove allosterically oriented the finger loop to point upward for core engagement with a GPCR. Furthermore, V2Rpp dissociation events suggest that its binding involves a sliding movement along the N-domain groove, during which its phosphorylated residues p3 and p4 transiently occupy the S5 site to facilitate repositioning of p5 into the S5 site, thereby triggering a zippering process of p1 to p3. The dynamic 62-residue βarr1 tail explored a vast conformational space, forming transient secondary structures, and could favorably anchor on the main body’s back side and within the central crest crevice. These findings elucidate key mechanistic steps underlying βarr1 activation.

## INTRODUCTION

G protein-coupled receptors (GPCRs) are integral membrane proteins that regulate many aspects of cell function and physiology and are targeted by more than one-third of all FDA-approved drugs (Pierce et al. 2002, Lefkowitz 2013, Hauser et al. 2017, Sriram and Insel 2018). When stimulated by an extracellular ligand, a GPCR interacts with its cognate G protein(s) on the cytoplasmic side of the receptor to initiate downstream signaling events (Weis and Kobilka 2018). Importantly, GPCRs can also activate distinct signaling pathways through the engagement of arrestins, particularly non-visual β-arrestins (βarr1 and βarr2, collectively βarr) (Peterson and Luttrell 2017). βarr are recruited by hundreds of different GPCRs to desensitize G protein signaling and to promote clathrin-mediated receptor endocytosis (Gurevich and Gurevich 2019). By scaffolding kinases and other molecules, receptor-activated βarr also facilitate diverse cellular signaling pathways distinct from those initiated by G proteins (Lefkowitz and Shenoy 2005, Shenoy and Lefkowitz 2011, Kahsai et al. 2018, Smith et al. 2018, Ahn et al. 2020). Unraveling the intricacies of βarr-mediated signaling to understand the selective activation of specific downstream pathways holds profound therapeutic implications (Shenoy and Lefkowitz 2011, Xu and Shao 2022).

Arrestins are mainly composed of two symmetrical and structured domains, known as the N and C domains and collectively referred to as the “main body” in this study (**Fig. 1A**). The loops that intersect these domains form what is referred to as the central crest region. In βarr, a groove in the N domain (N-domain groove) secures βarr’s own C-terminal tail (βarr tail) in its autoinhibited position in the basal state (Han et al. 2001, Asher et al. 2022). Specifically, the middle segment of the βarr tail, previously defined as residues 384-396 in βarr1 (Asher et al. 2022), adopts an extended β-strand conformation within the groove. In the active state of βarr, the phosphorylated receptor C-terminal tail (Rp tail), enriched with negative charges, binds to the positively charged βarr N-domain groove, displacing the middle segment of the βarr tail. Variations of the phosphorylated residues in the Rp tails of different receptors have been shown to affect the functions of βarr, acting as “barcodes” that βarr decode to elicit distinct cellular signals (Kim et al. 2005, Tobin 2008).

**Figure 1.**
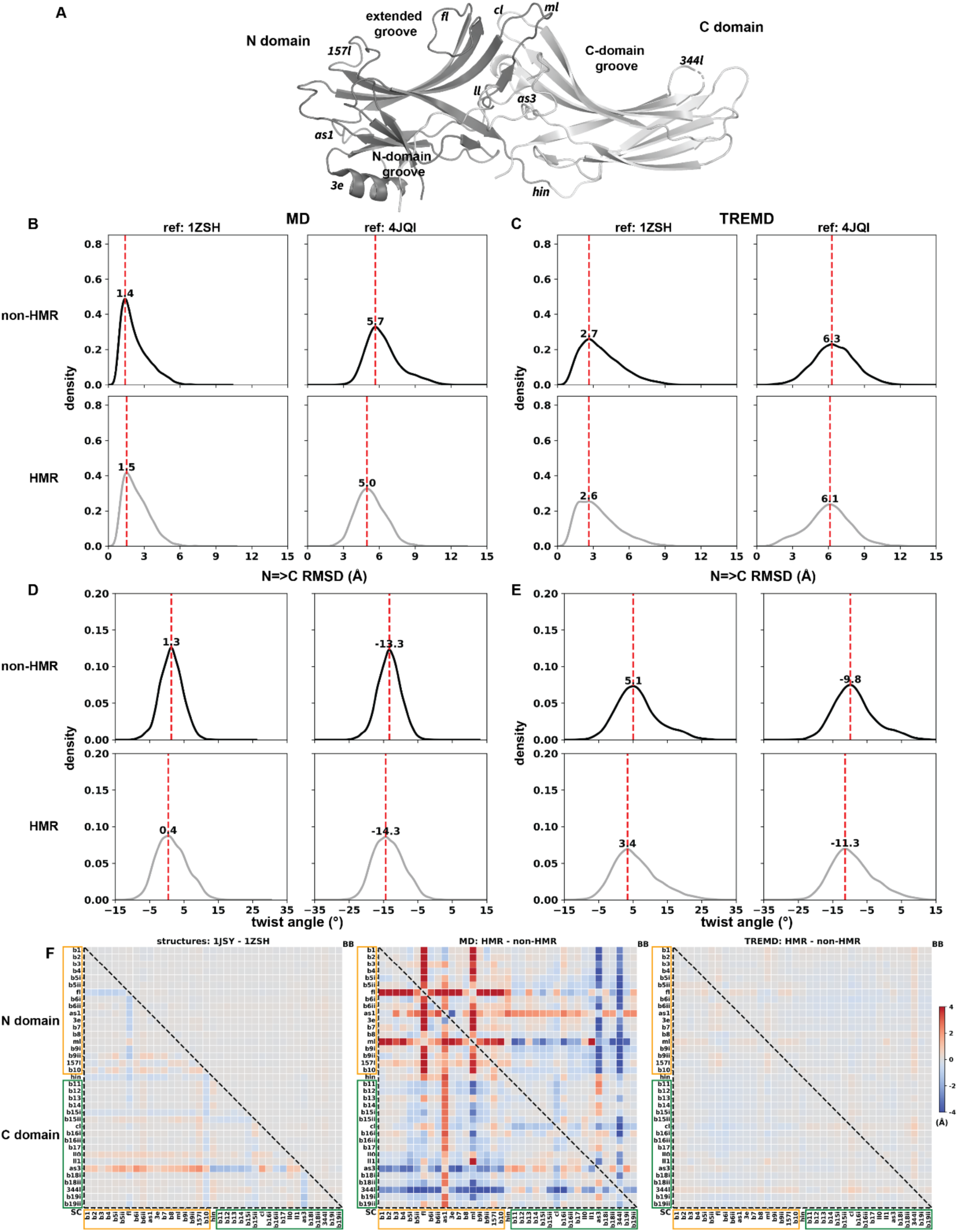
TREMD simulation of the basal state preserves the overall conformation while exploring a wider range of conformational space. (A) A βarr1 model in the basal state, highlighting key structural elements. See Table S5 for subsegment abbreviations. (B,C) “N⇒C” RMSD distributions from MD (B) and TREMD (C) simulations of the basal state, measured relative to 1ZSH (basal) and 4JQI (V2Rpp-bound active). TREMD shows broader sampling while maintaining proximity to the basal structure. (D,E) Twist angle distributions from MD (D) and TREMD (E) simulations using 1ZSH and 4JQI as references, further illustrating that TREMD maintains the basal-state conformation while enhancing conformational sampling. Red dotted lines indicate the peak values of the distributions in panels (B–E). (F) Heatmap comparison of subsegment rearrangements between hydrogen mass repartitioning (HMR) and non-HMR simulations assessed by PIA. The left panel compares two highly similar basal-state βarr1 structures (1JSY vs. 1ZSH), showing negligible backbone differences (top-right heatmap region) but some divergence in sidechain positions (lower-left region, enclosed by a black dotted triangle). The middle and right panels compare HMR and non-HMR conditions for MD and TREMD simulations, respectively. TREMD results are consistent across conditions even including sidechain positions, while MD simulations show divergence in flexible loop segments not captured by RMSD or twist angle measurements, indicating that the MD simulations had not converged. In all heatmaps, comparisons are labeled in the format “condition a – condition b” at the top of each panel; red pixels indicate subsegment distances that are larger in condition a, and blue pixels indicate those that are larger in condition b.

In addition to the Rp tail engagement, the intracellular portions of receptor transmembrane helices and cytoplasmic loops (receptor core) can also interact with the central crest loops of βarr (Shukla et al. 2014, Scheerer and Sommer 2017, Kahsai et al. 2018). Previous studies have demonstrated that receptor can engage βarr through either its Rp tail or both the receptor core and the Rp tail (Shukla et al. 2014). These modes are thought to confer distinct biological functions (Cahill et al. 2017). A recent study revealed that when the Rp tail of the Atypical Chemokine Receptor 2 (also referred to as the D6 receptor; D6Rpp) is bound to βarr2, the middle segment of the βarr2 tail transitions from a β-strand conformation when engaged with the N-domain groove into a helical conformation upon binding within the central crest pocket of βarr2. This structural adaptability has led to the middle segment of the βarr tail being referred to as the chameleon motif. Notably, occupation of the central crest pocket by this motif would be expected to prevent receptor core engagement (Maharana et al. 2024).

Elucidation of the structural rearrangements of βarr that occur upon receptor activation will facilitate the pharmacological targeting of specific regions, interactions, or molecular events. Compared to the basal state, crystal and cryo-electron microscopy structures of active βarr have revealed rearrangements in the central crest loops and rotation of the C domain relative to the N domain (Shukla et al. 2013, Kang et al. 2015, Staus et al. 2020, Maharana et al. 2024). However, the positions, conformations, and dynamics of the βarr tail with respect to the main body are unknown, as the βarr tail was either removed or largely unresolved in the experimentally determined structures (Gurevich et al. 2018, He et al. 2021). The βarr tail, characterized by its apparent flexibility, contains regions essential for binding clathrin and adaptin to mediate receptor endocytosis and may also be directly involved in scaffolding signaling kinases. Therefore, mechanistic insights into the functionally relevant conformational ensemble of the βarr tail in the context of the βarr main body are crucial (Gurevich and Gurevich 2015, Perry-Hauser et al. 2022). Furthermore, distinguishing the impact of the bound Rp tail from that of the βarr tail on the structural rearrangements of the main body is critical for understanding how distinct interactions within the N-domain groove shape the conformational landscape of βarr activation.

Considering the dynamic and at least partially unstructured nature of the βarr tail, molecular dynamics (MD) simulation is a suitable approach to sample and characterize its conformational ensemble. Nevertheless, the 62-residue-long βarr1 tail can in principle move extensively around the main body and may form transient secondary structures within a large three-dimensional space. Achieving adequate sampling of its conformational landscape remains beyond the timescale accessible to classical MD simulations. Addressing this limitation necessitates the applications of enhanced sampling techniques, such as temperature replica-exchange MD (TREMD) simulations (Sugita and Okamoto 1999, Rhee and Pande 2003), which can significantly improve sampling efficiency within a practical runtime. In TREMD, system configurations are periodically exchanged between two adjacent replicas according to a Metropolis acceptance criterion, facilitating the sampling of rare states at elevated temperatures (Metropolis et al. 1953, Sugita and Okamoto 1999, Qi et al. 2018). TREMD is a specialized implementation of the Metropolis-Hastings algorithm, employing Markov chain to ensure that each replica samples from the correct canonical ensemble while maintaining reversibility (Hastings 1970). When a temperature ladder is constructed with an appropriate temperature increment, i.e., comparable to the system’s potential energy barriers, the sampling efficiency of TREMD is greatly enhanced compared to classical MD simulations conducted at a fixed temperature (typically room or physiological temperature) (Kofke 2002, Nymeyer 2008). Previous studies showed TREMD performed on various systems can achieve sampling efficiencies from 5-fold to a few orders of magnitude faster than classical MD simulations (Sanbonmatsu and Garcia 2002, Rosta and Hummer 2009).

In this study, we used TREMD simulations to investigate the activation mechanism of βarr1 by characterizing and comparing the conformational dynamics of both the basal and active states and identified novel and likely functionally relevant conformational ensembles of βarr1 tail in the active state.

## RESULTS

To comprehensively investigate the conformational dynamics and transitions between the basal and active states of βarr1, we used the crystal structures of arrestins as templates to construct full-length models for both states, including modeling the regions missing in the structures. In particular, since the βarr1 tail was only partially resolved in the basal state and completely unresolved in the active state, we adopted both homology and ab initio modeling approaches. For the active state, we modeled βarr1 bound to V2Rpp, a synthetic phosphorylated peptide mimetic of the human V2 vasopressin receptor tail, for which we also modeled several residues missing from the crystal structure (see **Methods** for model construction details). To enhance conformational sampling efficiency, in addition to classical MD (hereafter referred to as “MD”), we also conducted TREMD simulations for both states (**Table S1**) (See Methods for the simulation protocols and **Supporting Information** and **Figs. S1 and S2** for the evaluation of the sampling efficiency). These simulations generated a very large data set, prompting us to adapt as well as to develop clustering, geometric, and energetic analytical methods to elucidate the relevant conformational states and dynamics, as described below and in **Methods**.

### TREMD simulation of the βarr1 basal state samples a broad conformational space but maintains the overall state

Using both the MD and TREMD simulation results of the basal state, we first compared the sampling extent of these two simulation approaches to evaluate the robustness and reliability of our simulation protocols. Specifically, we used the crystal structures of the basal (PDB 1ZSH) and active (PDB 4JQI) states of βarr1 as references and assessed two geometric observables, namely the root mean square deviation (RMSD) and the twist angle.

In the RMSD analysis, we first compared three alignment-calculation schemes (see **Methods** for the definitions of the residue ranges used in alignment and RMSD calculations). Note that we included only the structured segments of the indicated domains in these schemes, i.e., excluding the loop regions. Specifically, we tested: (1) aligning the entire main body to calculate its RMSD (termed the main-body scheme), (2) aligning the N domain to calculate the C-domain RMSD (N=>C), and (3) aligning the C domain to calculate the N-domain RMSD (C=>N). While all three schemes displayed similar trends for the basal-state MD simulations (**Fig. S3**), the latter two schemes revealed more clearly the divergence between the basal and active states, offering a significantly larger dynamic range compared to the main body scheme. For example, using the main body scheme, the peak value of the RMSD distribution for the basal-state MD simulation results was 2.2 Å with respect to the active-state structure 4JQI, which is 1.3 Å larger than that with respect to the inactive-state structure 1ZSH, a difference not significantly larger than the RMSD fluctuations observed in MD simulations. In contrast, using the N=>C scheme, the peak RMSD value with respect to 4JQI increased markedly to 6.0 Å, which is 4.5 Å larger than that with respect to 1ZSH. Consequently, we focused on the N=>C scheme for the subsequent comparisons, as it also corresponds well with the twist angle definition described below.

Our analysis revealed that the N=>C RMSD distributions peaked at 1.5 Å and 2.3 Å for the MD and TREMD simulations of the basal state, respectively, when using the basal 1ZSH structure as the reference (**Fig. 1B,C**). When the active 4JQI structure was used as the reference, the peak values were 6.0 Å and 6.7 Å for MD and TREMD, respectively. Although the peak values from the two types of simulations were similar, the TREMD simulations exhibited broader distributions (**Figs. 1B,C, S3, and S4**). These results suggest that the TREMD simulations enabled a broader exploration of conformational space than the MD simulations, while still reliably retaining the βarr1 conformation in the basal state.

The twist angle has been widely used as a metric to describe the extent of the relative rotation between the N and C domains of arrestins during the transition from the basal to active states. Because it has been vaguely defined previously, we developed a detailed definition of the structural elements and an algorithm for calculating this twist angle. Briefly, after aligning any two arrestin structures or models by their N domains, with one serving as the reference, the rotation angle is derived from the rotation matrix of the corresponding C domains. The twist angle is defined as the projection of this rotation angle onto the plane perpendicular to the axis from the center of mass (COM) of the N domain to the COM of the C domain in the reference structure (see **Supporting Information**, **Fig. S5**, **Table S2** for a detailed description and evaluation).

Our results showed that the twist angle distribution for the basal state, using the basal 1ZSH structure as the reference, peaked at 0.9° and 3.9° for the MD and TREMD simulations, respectively. When using the active 4JQI structure as the reference, the twist angle distributions peaked at -15.5° and -12.7° for the MD and TREMD simulations, respectively, demonstrating significant differences from the active structure (**Fig. 1D,E**). Consistent with the RMSD analysis, the broader twist angle distributions in TREMD compared to MD further confirmed that the TREMD simulations explored a broader conformational space while preserving the overall βarr1 conformation in the basal state (**Fig. 1D,E**).

In summary, TREMD simulations broadened conformational sampling while preserving the basal βarr1 conformation. In addition, as detailed in **Methods**, hydrogen mass repartitioning (HMR) was used to accelerate simulations, and comparison with non-HMR results confirmed it did not affect TREMD sampling.

### The main body of active V2Rpp-bound βarr1 transitions toward the basal state in the absence of Fab30

Using an active βarr1 structure bound with V2Rpp (PDB 4JQI) as the template for the main body of βarr1, we employed both homology and *ab initio* modeling approaches implemented in Modeller and Rosetta to build an active state model of βarr1 (see **Methods**). Unlike the structure 4JQI, our model excluded the Fab30 antibody fragment, which was used to stabilize the active state of βarr1 in the structural studies (Shukla et al. 2013). The resulting model includes an *ab initio-*modeled βarr1 C-terminal tail that was predominantly unwound. Starting from this model, we carried out both MD and TREMD simulations to investigate the conformations and dynamics of the main body of βarr1, the bound V2Rpp, and the βarr1 tail.

Visual inspection revealed that the main body of βarr1 bound with V2Rpp explored noticeably larger conformational space in TREMD compared to MD (**Fig. 2A**). We quantitively evaluated the main body conformations using the N=>C RMSD and twist angle, with both the structures 1ZSH (basal) and 4JQI (active) as references. In the MD simulations, the N=>C RMSD distributions peaked at 3.0 Å with respect to the starting model based on 4JQI, indicating a noticeable deviation from the Fab30-stabilized state. This value is comparable to the RMSD relative to the basal-state structure 1ZSH (3.5 Å), suggesting that the main body adopted an intermediate conformation between the two states (**Fig. 2B**). The twist angle values of 7.0° and - 10.8° with respect to 1ZSH and 4JQI, respectively, also supported this interpretation (**Fig. 2D**). These deviations were consistent with a potential transition towards the basal state.

**Figure 2.**
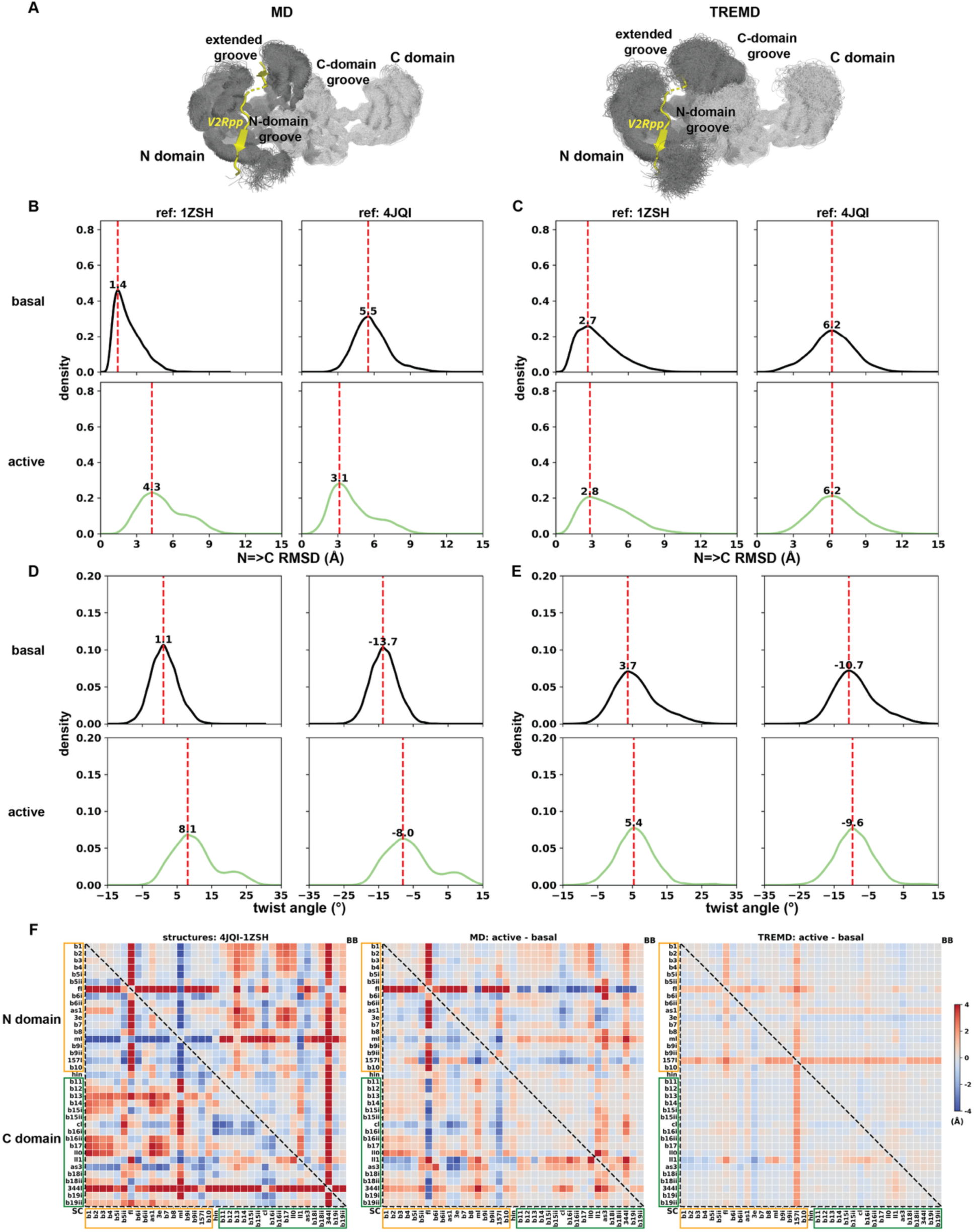
The main body of V2Rpp-bound βarr1 transitions toward the basal state in the absence of Fab30. (A) Superimposed βarr1 models of the V2Rpp-bound active state (without Fab30) from the MD (left) and TREMD (right) simulations, highlighting the significantly expanded conformational space of the main body explored by TREMD. (B,C) “N⇒C” RMSD distributions from MD (B) and TREMD (C) simulations of the V2Rpp-bound state, measured relative to 1ZSH (basal) and 4JQI (Fab30-stabilized active). TREMD shows broader sampling and a more pronounced shift away from the starting conformation. For comparison, combined RMSD distributions of basal-state HMR and non-HMR simulations are also shown. (D,E) Twist angle distributions from MD (D) and TREMD (E) simulations using 1ZSH and 4JQI as references, also including the combined basal-state simulations for comparison. TREMD results show a further shift toward the basal-state conformation. Red dotted lines indicate the peak values of the distributions in panels (B– E). (F) Heatmap comparison of subsegment rearrangements between the V2Rpp-bound and basal states assessed by PIA. The left panel compares the crystal structures of 4JQI and 1ZSH, showing rearrangements in both interdomain orientation and loop positions. The middle and right panels compare V2Rpp-bound MD and TREMD simulations, respectively, with the combined basal-state simulations. While the V2Rpp-bound TREMD simulations show an overall shift toward the basal state, the finger and 157 loops still exhibit marked divergence from their basal conformations. Color scheme and presentation follow the description in the Figure 1 legend.

The trend of transition towards the basal state became more pronounced in the TREMD simulations, where the N=>C RMSD distributions peaked at 2.8 Å and 6.7 Å with respect to 1ZSH and 4JQI, respectively (**Fig. 2C**). Moreover, the twist angle peaked at 4.6° with 1ZSH and -11.9° with 4JQI, closer to the basal-state reference and farther from the Fab30-stabilized conformation, consistent with a greater shift toward the basal state compared to the MD simulations (**Fig. 2E**). The more pronounced deviation of TREMD results from the starting conformation suggest that TREMD simulations overcome energy barriers that were difficult to cross in MD.

To further characterize conformational changes between the basal and active states beyond the relative movements of the N and C domains at the domain level, we adapted our previously developed Protein Interaction Analyzer (PIA) (Stolzenberg et al. 2016, Michino et al. 2017, Lee et al. 2023) for this study. By dividing the protein into subsegments and analyzing changes in their pairwise distances between two different states, PIA enables characterization of structural rearrangements at the structural motif level (Stolzenberg et al. 2016, Michino et al. 2017).

Notably, this approach is superposition independent and does not rely on any reference structures. For this study, the subsegments were defined by integrating those previously described (Hirsch et al. 1999, Gurevich et al. 2018), with minor adjustments informed by visual inspection of experimentally determined structures, and by dividing long β-strands into shorter subsegments to enhance resolution (see **Methods** for residue ranges of each subsegment). As a control analysis, we compared the simulation results of the basal state with and without applying the HMR technique. Consistent with the RMSD and twist angle analysis, the TREMD simulations showed minimal differences between the two conditions in the PIA analysis. In contrast, the corresponding MD simulations exhibited substantial divergence in certain loop regions in the PIA (**Fig. 1F**). These differences were not captured by the RMSD and twist angle metrics described above, which only assess the structured regions of the main body (**Fig. 1B,D**). These loop-level differences suggest that, at the current simulation length (∼4 µs), the MD trajectories had not fully converged, whereas the TREMD simulations do converge at this timescale.

Our PIA analysis comparing the MD simulation results of the basal and V2Rpp-bound βarr1 states revealed similar differences as those between the structures 4JQI and 1ZSH. Specifically, in the structured regions of the main body, the β11-β17 region of the C domain generally exhibited longer distances to the β1-β4 and the arrestin switch I (ASw1)-β7 regions of the N domain in the V2Rpp-bound state, consistent with the relative rotations between the C and N domains observed in the two states (**Fig. 2F**). The most significant rearrangements in the active state occurred in the finger and middle loops: the finger loop showed increased distances to the N domain and shorter distances to the C domain, while the middle loop exhibited shorter distances to the N domain and longer distances to the C domain.

Strikingly, however, except for the changes in the finger loop, these rearrangements were significantly reduced in the TREMD simulations comparing the two states. In particular, the rearrangements in the β11-β17 region of the C domain almost disappeared entirely, consistent with the aforementioned small twist angle of the V2Rpp-bound state with respect to the basal 1ZSH structure, but a large twist angle when using 4JQI as the reference. Interestingly, the so-called 157 loop showed increased distances to both domains in the V2Rpp-bound state during TREMD simulations, which was not observed in the MD simulations (**Fig. 2F**).

Taken together, our three analyses indicated that during the TREMD simulation of the V2Rpp-bound state without Fab30, the main body of βarr1 exhibited a strong tendency to transition back toward the basal state, consistent with the need for the Fab30 to stabilize the active state in the structural studies.

### Allosteric modulation of the finger loop by V2Rpp binding in the N-domain groove

The major difference between the basal and V2Rpp-bound states arises from the occupancy of the N-domain groove: in the basal state, it is occupied by the middle segment of the βarr1 tail, whereas in the V2Rpp-bound state, it is occupied by V2Rpp, which additionally extends into the extended groove of the N domain. V2Rpp contains eight phosphorylated Ser and Thr residues designated p1 through p8. The p1-p3 region binds to the extended groove, the p5-p8 region interacts with the N-domain groove, and p4 is situated between these two grooves and exposed to the aqueous environment. While p5 to p8 collectively carry eight negative charges, the corresponding residues of the βarr1 tail occupying the same space in the N-domain groove carry only two. In addition to their sequence differences, V2Rpp and the βarr1 tail bind in opposite orientations in the N-domain groove, resulting in different backbone interactions with the main body (**Fig. 3B,C**). However, the similarity of the RMSD and twist angle between the basal and V2Rpp-bound states observed in our TREMD simulations suggested that these difference in binding have limited impact on the structured portions of the main body.

**Figure 3.**
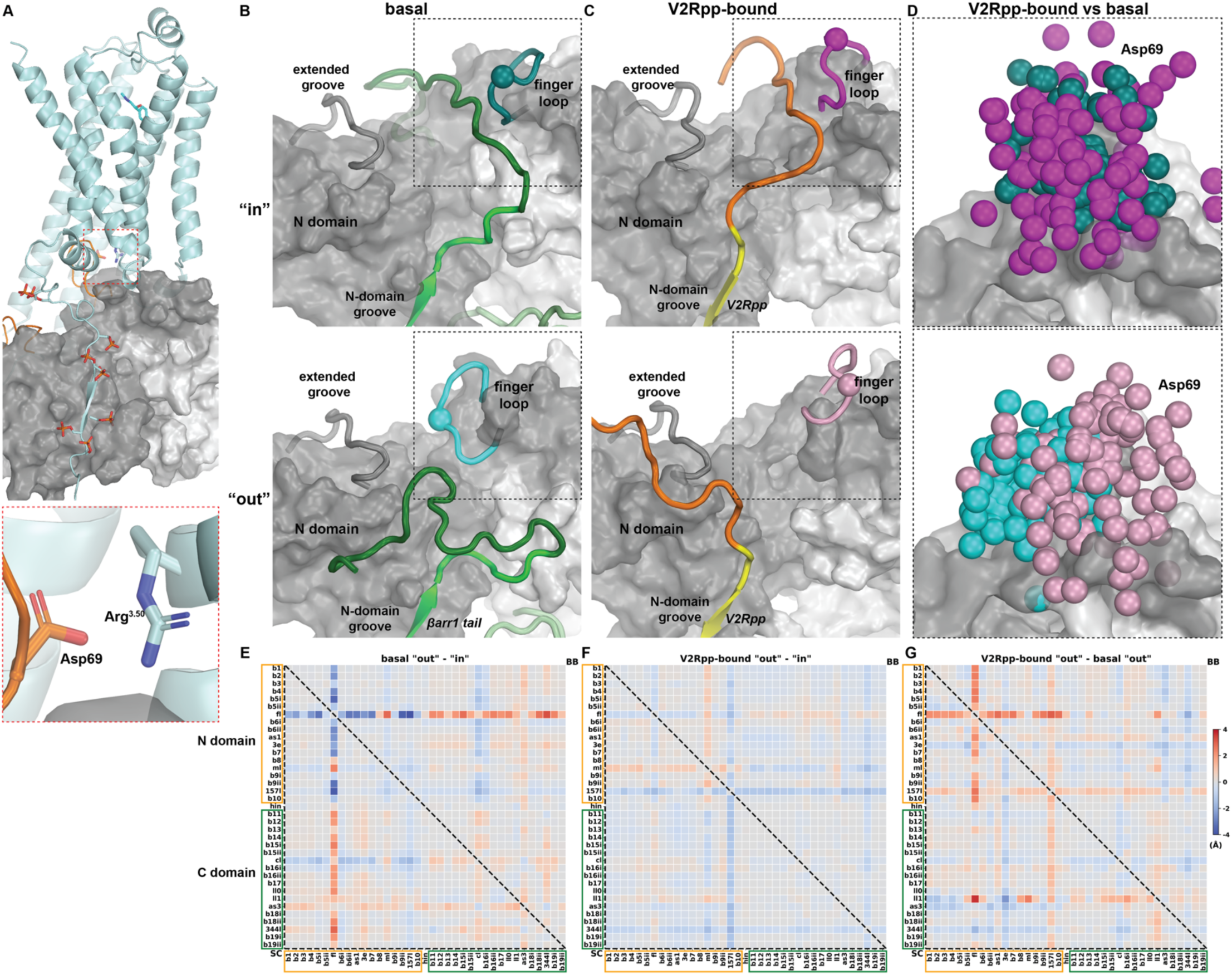
V2Rpp binding in the N-domain groove allosterically modulates the finger loop conformation. (A) The structure of βarr1 in complex with the β1-adrenergic receptor (B1AR) highlights the upward orientation of the finger loop in the active state. B1AR is shown as a cyan ribbon, and the finger loop is colored orange. The upward orientation enables an ionic interaction between βarr1 Asp69 and B1AR Arg^3.50^, highlighted by the red-dotted box. (B,C) Representative snapshots from TREMD simulations of the basal state (B) and V2Rpp-bound state (C), grouped into “in” (upper panels) and “out” (lower panels) substates based on whether the βarr1 tail (basal) or V2Rpp (active) occupies the extended groove. (D) Overlay of finger loop conformations from the “in” and “out” substates of both states, with the Cα atoms of Asp69 shown as spheres colored to match the corresponding structures in (B) and (C). The V2Rpp-bound state consistently shows an upward finger loop conformation regardless of extended groove occupancy, whereas in the basal state, the finger loop collapses into the extended groove when the βarr1 tail does not occupy it. (E,F) PIA comparisons between the “in” and “out” substates in the basal (E) and V2Rpp-bound (F) simulations reveal a significant rearrangement of the finger loop only in the basal state. (G) PIA comparison between the “out” substates of the basal and V2Rpp-bound simulations further demonstrates that V2Rpp binding in the N-domain groove promotes an upward orientation of the finger loop. Subsegment definitions follow Table S5. Color scheme and labeling follow the description in the Figure 1 legend.

Therefore, we sought to identify other structural features that distinguish these two states. Based on the PIA analysis results, we focused on the finger loop, which forms one side of the extended groove. The finger loop plays a critical role in facilitating the core engagement between arrestin and GPCRs (Zhou et al. 2017, Lee et al. 2020, Asher et al. 2022). In the basal βarr1 structure 1ZSH, the finger loop is collapsed into the extended groove, hindering its interaction with the receptor core. Conversely, in the V2Rpp-bound βarr1 structure 4JQI, the finger loop adopts an upward orientation, positioning it for effective receptor engagement, such as that demonstrated by the structure of βarr1 in complex with the β1-adrenergic receptor (PDB 6TKO, **Fig. 3A**).

In the TREMD simulations of the basal state, the distant segment of the βarr1 tail occasionally entered the extended groove (referred to as the “in” substate), occurring in 1.6% of the frames. This entry was coupled with a flipping rearrangement of the finger loop, which opened the extended groove. As a result, the finger loop adopted a conformation drastically different from that observed in the basal state structure 1ZSH. Conversely, when the βarr1 tail did not occupy the extended groove (“out” substate), the finger loop largely collapsed into the groove in the simulations, consistent with structure 1ZSH (**Fig. 3B**). This variation in the finger loop orientations was further reflected in a PIA analysis comparing the “out” and “in” substates, which showed that in the “out” substate, the finger loop was positioned closer to the N-domain segments and farther from the C-domain segments (**Fig. 3E**).

In our TREMD simulations of the V2Rpp-bound state, we observed a dynamic behavior of V2Rpp that could not be inferred from the static structure 4JQI. Specifically, the first half of V2Rpp, including the p1 to p3 region, could somewhat stably occupy the extended groove (“in” substate) but also frequently dissociated from it (“out” substate). In contrast, the second half remained associated with the N-domain groove, maintaining its interactions unchanged in the vast majority of the simulation frames (see the next section for an analysis of other scenarios). Unlike the significant rearrangement of the finger loop observed in the basal state depending on whether the βarr1 tail entered the extended groove, the finger loop in the V2Rpp-bound state consistently remained pointed upwards, regardless of whether V2Rpp was bound in the extended groove (**Fig. 3C**), as reflected in a PIA analysis between the “out” and “in” substates (**Fig. 3F**). Furthermore, by comparing the finger loop orientations across the basal and V2Rpp-bound substates, we found that the finger loop in the V2Rpp-bound “out” substate was significantly more open than in the basal “out” substate (**Fig. 3D,G**).

The observations of the basal “in” substate and V2Rpp-bound “out” substate, both diverging from the corresponding crystal structures, provided a comprehensive picture of the dynamics of the bound βarr1 tail and V2Rpp, as well as the modulation of the grooves. Together, our results suggested a correlation between V2Rpp binding in the N-domain groove and the finger loop conformation, while βarr1 tail binding did not exhibit such a long-range impact. Thus, we propose that V2Rpp allosterically modulates the finger loop, orienting it upwards for core engagement with GPCRs, through its binding within the N-domain groove. In contrast, its binding in the extended groove does not appear to be critical for this effect.

### The binding and dissociation profiles of the phosphorylated residues of V2Rpp

Substantial evidence has emerged in support of the barcode hypothesis, which proposed that the distinct phosphorylation patterns of Rp tail residues trigger different arrestin-dependent signaling outcomes (Kim et al. 2005, Tobin 2008). Previous computational and experimental studies have explored and identified key phosphorylated residues in the Rp tails (Latorraca et al. 2018, He et al. 2021, Maharana et al. 2023). However, critical mechanistic questions remain unanswered, including the potentially distinct functional roles and binding mechanisms of the Rp tail in the N domain versus the extended grooves. This gap in understanding is partly due to the absence of studies, to our knowledge, that have elucidated the dissociation process of the bound Rp tail, such as V2Rpp, from βarr1.

For the eight phosphorylated residues in V2Rpp (p1 to p8), their corresponding binding sites revealed in the V2Rpp-bound βarr1 structure (PDB: 4JQI) are referred to as the S1 through S8 sites in this study (**Fig. 4A**). Interestingly, none of these eight sites is formed in the basal state, despite the presence of negatively charged Glu389 and Asp390 from the βarr1 tail anchored in the N-domain groove, as large rearrangements of the finger, middle, and lariat loops during arrestin activation are required for their formation. To investigate the conformational dynamics explored by V2Rpp in the MD and TREMD simulations of the V2Rpp-bound βarr1 state, we measured the deviations of the phosphate groups of p1-p8 in the simulations from their original positions in 4JQI. We found that V2Rpp in TREMD explored significantly broader conformational space than in MD (**Figs. 4B**). Specifically, in TREMD, as described above, the p1-p3 region could frequently and completely dissociate from the extended groove (deviation > 10 Å), while p5-p8 region exhibited deviations of up to 10 Å from their original bound positions (**Fig. 4C**). In contrast, during the MD simulations, other than p4, the phosphorylated residues rarely deviated beyond 5 Å.

**Figure 4.**
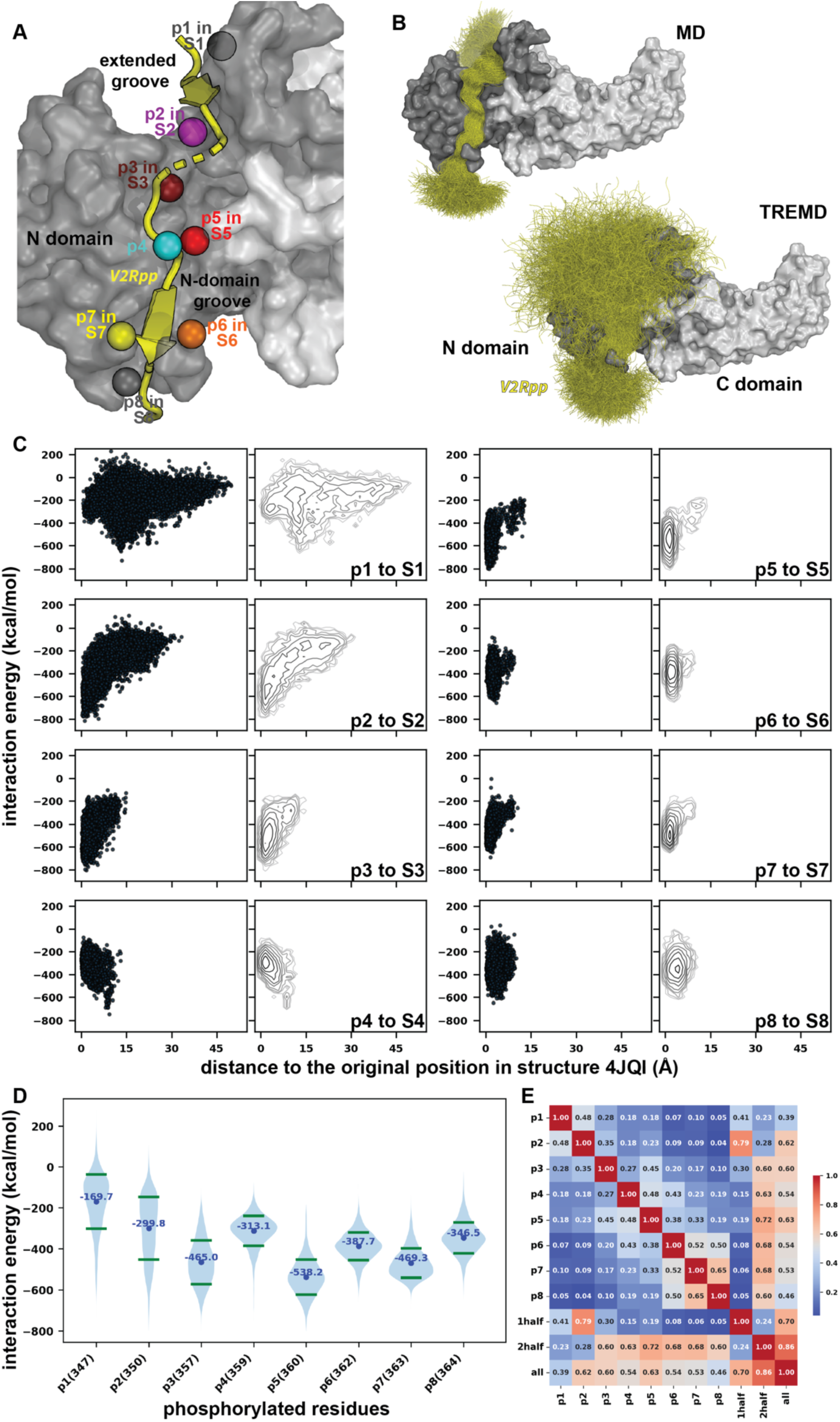
p5 exhibits the strongest interaction with βarr1. (A) The structure of V2Rpp bound to βarr1 (PDB: 4JQI), showing the eight phosphorylated residues (p1–p8) and their corresponding binding sites (S1–S8) in the N-domain and extended grooves. (B) Distributions of the bound V2Rpp conformations in MD (left) and TREMD (right) simulations, highlighting broader conformational sampling, including partial dissociation of V2Rpp, in TREMD. (C) Distributions of positional deviation versus interaction energy for each phosphorylated residue (p1–p8) from TREMD simulations. For each residue, the left panel shows a scatter plot, and the right panel shows a contour plot of the same data. Positional deviation is measured for the phosphorus atom of the phosphate group relative to its crystallographic position in 4JQI (i.e., the distance of p1 to the S1 site (“p1 to S1”), “p2 to S2”, etc.), and interaction energy reflects the entire phosphorylated residue’s interaction with βarr1. Darker contours indicate regions of higher density. p5 showed both the smallest deviation and the most favorable interaction energy. (D) Average interaction energies of individual phosphorylated residues with βarr1, showing that p5 binds most strongly, followed by p7 and p3. (E) Correlation matrix of interaction energies among phosphorylated residues and between residue-level and segment-level energies, revealing coordinated binding behavior (see text) and highlighting the most influential residues in each half of the peptide: p2 in the first half (p1-p2), and p5 in the second half (p3-p8).

To estimate the binding strengths of each phosphorylated residue, we calculated the interaction energies between each residue and the entire βarr1 in the simulations (see **Methods**). In TREMD, the results showed that p5 exhibited the strongest interactions with βarr1, with an average interaction energy ∼70 kcal/mol more favorable compared to the other phosphorylated residues (**Fig. 4C,D**). p5 was followed by p7 and p3, which showed comparable interaction energies. Among the remaining phosphorylated residues, p6 and p8 displayed significantly stronger binding than p4 and p2, while p1 was the weakest binder (**Fig. 4C,D**). This binding strength trend of p1-p8 largely aligned with the extents of the deviations observed for these residues from their original positions (see above), except for p4 (**Fig. 4C**). The position of p4 is restrained by its two strong neighboring binders, p5 and p3, and, consequently, p4 appeared to have even smaller deviations than p3, despite not forming strong interactions with βarr1 itself. In contrast, in the MD simulations, p2 appeared to be the strongest binder, while the binding strength trends for other residues varied across different MD trajectories. This variability is likely due to the residues being trapped in different local energy minima on the energy surface, reflecting inadequate sampling in the MD simulations.

Building on these observations, we conducted a correlation analysis of the interaction energies for individual phosphorylated residues and different segments of V2Rpp to further explore the interplay between binding strengths and coordination among residues. The PxPP motif, recently proposed as a key element in a common ’lock-and-key’ activation mechanism of βarr by GPCRs (Maharana et al. 2023), highlights potential coordination among p5 to p7 (Maharana et al. 2023). Our results revealed a coordinated binding relationship between p7 and p8, and to a lesser extent, between these residues and p6. Interestingly, p5 binding exhibited a stronger correlation with p4 than with p6, which was unexpected given the proposed PxPP motif suggesting potential coordination among p5 to p7 (**Fig. 4E**). Furthermore, the interaction energies of p2 and p5 were most strongly correlated with those of the first and second halves of V2Rpp, respectively, with p5 showing the highest correlation with the interaction energy of the entire V2Rpp (**Fig. 4E**). These findings suggested a complex interplay among phosphorylated residues in modulating V2Rpp binding.

Notably, the identification of p5, the first P in the proposed PXPP motif, as the strongest binder in TREMD is consistent with a previous crystallography study. In that study, dephosphorylation of p5 was shown to have the most notable impact on the bound V2Rpp conformation, not only disrupting the local interaction network but also propagating to a loop on the opposite side of the βarr1 main body from the N-domain groove (PDB 7DFA) (He et al. 2021).

Interestingly, our energetic calculations, combined with the deviation analysis, suggested that the original binding sites of p1 and p4, i.e., the S1 and S4 sites, in the structure 4JQI were not their most favored binding spots in the grooves (**Fig. 4C**). Thus, these two residues could potentially occupy other sites or intermediate binding positions before transitioning to their final configuration (see below), which was then stabilized by neighboring stronger-binding phosphorylated residues.

### Binding of V2Rpp to βarr1 may involve an initial sliding followed by a zippering motion

A critical aspect of the recognition between p1-p8 and βarr1 is the ionic interactions between the positively charged phosphate groups of these phosphorylated Ser and Thr residues, which share extremely similar physiochemical properties, and the negatively charged sidechains of Arg and Lys residues located in or surrounding the grooves, forming the S1 to S8 sites. The arrangement and coordination of these ionic interactions are thought to form the structural basis of the barcode hypothesis.

To further characterize binding strength and coordination of these phosphorylated residues, we defined binary interaction states for each residue after identifying all their interacting Arg and Lys residues in the 4JQI structure (termed as interaction partners, see Methods for the list). A state of "1" indicated that the residue retained at least one ionic interaction with their original interaction partners, while "0" indicated that all such ionic interactions were lost. Note that p4 does not directly interact with any Arg and Lys residue in 4JQI in which the closest Arg and Lys to p4 is Lys294, which we arbitrarily defined as its interaction partner. Consequently, the interaction pattern for p1-p8 in the 4JQI structure is 1-1-1-0-1-1-1-1, signifying that all phosphorylated residues retain at least one original ionic interaction, except for P4.

We then quantified and analyzed the occurrence of all possible interaction patterns between p1-p8 and βarr1 in the TREMD simulations. The most frequent interaction patterns in our simulations were <u>0</u>-1-1-0-1-1-1-1 (29.0%) and <u>0</u>-<u>0</u>-1-0-1-1-1-1 (28.1%), highlighting relatively weak binding of p1 and p2, consistent with our interaction energy and deviation calculations. Interestingly, p4 exhibited a significant probability of interacting with Lys294 in the simulations, with the patterns 0-0-1-<u>1</u>-1-1-1-1 (7.7%) and 0-1-1-<u>1</u>-1-1-1-1 (5.5%) showing even higher probabilities than the original 1-1-1-<u>0</u>-1-1-1-1 (5.4%) configuration. Among the seven most frequently observed patterns, which accounted for over 80% of the simulation frames, p5 through p8 were all in their bound state (“1”). This finding indicated that the portion of V2Rpp interacting with the N-domain groove had a stronger binding than the first half of the peptide. Further analysis of the 20 most frequent patterns, comprising more than 94% of the frames, showed that both p5 and p7 were consistently bound in their original S5 and S7 sites, respectively, whereas p3 exhibited a significant likelihood of losing all its original ionic interactions. Interestingly, this observation diverged from the results of interaction energy calculations, which suggested that p3 and p7 have comparable binding strengths. Thus, interaction energies for individual phosphorylated residues could not fully capture the dynamic coordination of phosphorylated residues.

When p5 dissociated from the S5 site, which occurred in approximately 1.6% of TREMD frames, we observed that V2Rpp did not detach from the grooves but instead slid in the N-domain groove toward the direction of the S8 site. Specifically, we identified two clusters of V2Rpp configurations: one where p4 slid into the S5 site in which the phosphate was coordinated by Lys11, Arg25, and Lys294 of βarr1 (termed the “p4-in-S5” state); and another where p3 occupied this site (“p3-in-S5”). In the S5 site, p4 exhibited more favorable interaction energy with βarr1 compared to in its original site (as shown by the cyan points in **Fig. 5A**), while p5 transitioned to a relatively high-energy state in an exposed location between the S5 and S6 sites (termed as the S5’ site, cyan points in **Fig. 5C**). Interestingly, when p3 occupied the S5 site, p4 was at the S5’ site, while p5 itself was approximately at the S6 site (**Fig. 5B,C**). Notably, in the rare occurrence of the 0-0-0-0-0-0-0-0 pattern, characterized by the loss of all original ionic interactions of the phosphorylated residues, all the frames corresponded to either the “p3-in-S5” or “p4-in-S5” state.

**Figure 5.**
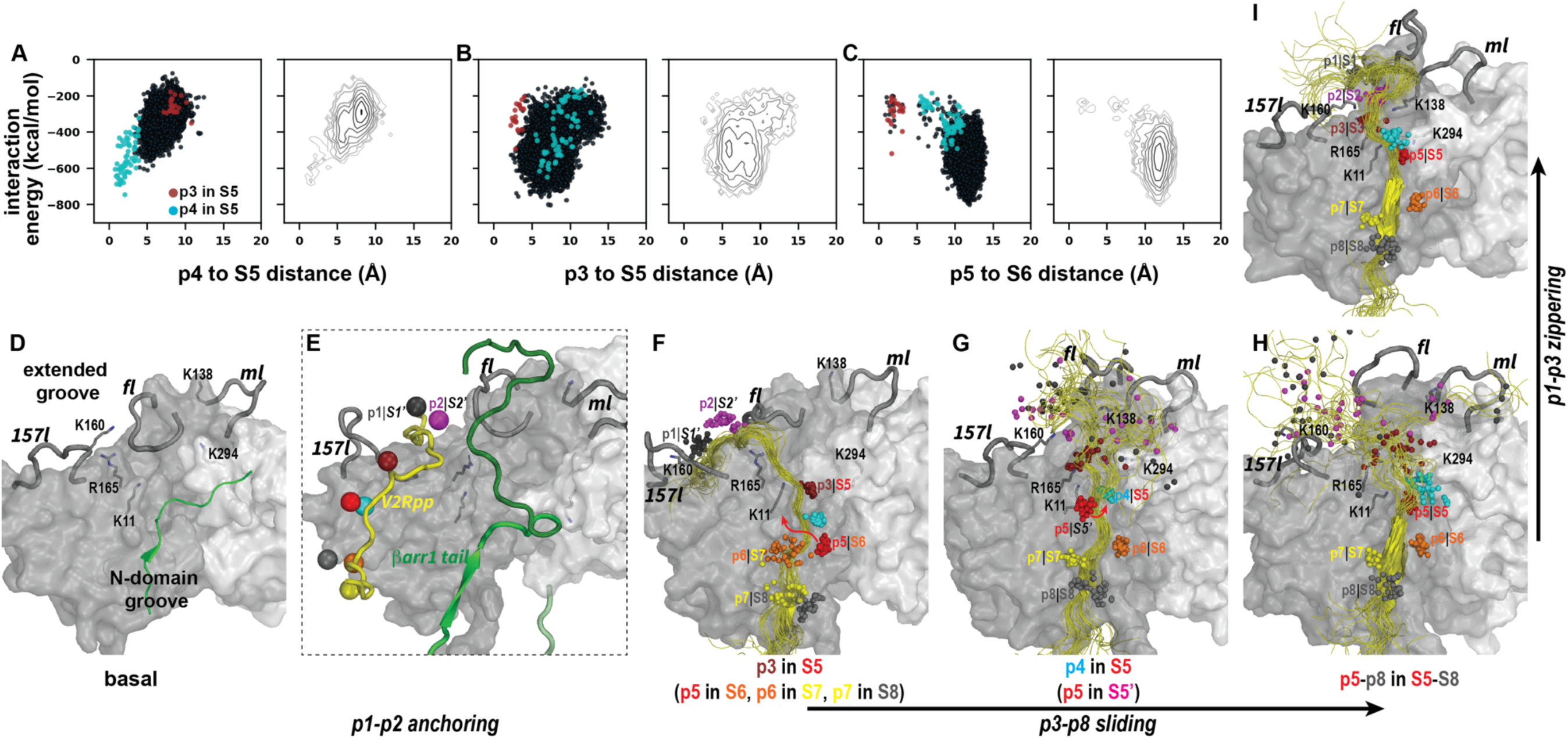
Binding of V2Rpp to βarr1 involves both sliding and zippering motions. (A–C) Distributions of distance versus interaction energy for individual phosphorylated residues from TREMD simulations. Each plot shows the distance between the phosphate phosphorus atom and its alternative binding site, plotted against the interaction energy of the entire phosphorylated residue with βarr1. Panel A shows the distance from p4 to the S5 site, panel B from p3 to S5, and panel C from p5 to S6. The format is consistent with Figure 4C. (D) The basal-state structure of βarr1 (PDB 1ZSH), showing occupation of the N-domain groove by the βarr1 tail (green) and collapse of the finger loop into the extended groove, corresponding to the basal “out” substate shown in Fig. 3B. (E) Zoomed-in view of the grooves showing a proposed intermediate state in which the first half of V2Rpp (p1-p2) initially anchors in alternative sites (S1′ and S2′) in the extended groove, with the finger loop occupying the groove. This anchoring would position the second half of V2Rpp (p3-p8) to compete with the βarr1 tail for binding to the N-domain groove. (F) Representative frames from TREMD simulations showing the “p3-in-S5” state, in which p1 and p2 remain engaged with the S1′ and S2′ sites, and the finger loop is collapsed into the extended groove. While p3 binds energetically favorably to S5, this is not its most favorable position (as shown in panel B). Compared to the basal state, Lys294 is repositioned to participate in S5 formation. (G) Representative frames showing the “p4-in-S5” state, where p4 occupies its most favorable position observed in TREMD (see panel A). Compared to panel F, the finger and middle loops have rearranged to open the extended groove, and p1-p3 have dissociated from their earlier transient positions. (H) Representative frames showing p5-p8 docked in the final S5-S8 sites, while p1-p3 remain dynamic and intermittently occupy the extended groove. The finger loop adopts an upward orientation, and Lys65, Lys138, and Lys160 shift into their active-state positions to form the S1, S2, and S3 sites. (I) Schematic representation of the proposed zippering mechanism, in which p1–p3 progressively engage the extended groove following anchoring of p5–p8 in the N-domain groove, thereby establishing the S1–S3 sites. These results support a stepwise V2Rpp binding mechanism involving an initial sliding phase followed by progressive stabilization through zippering interactions.

A further examination of the “p4-in-S5” state revealed that the middle loop, which contributed Lys138 to the S2 and S3 sites, swung away toward the C domain (**Fig. 5G**). In the “p3-in-S5” state, the conformational changes of both the middle and “157” loops, caused both Lys138 and Lys160 to reorient away from these two sites, adopting their positions in the basal state (**Fig. 5D**). In coordination with these movements, the finger loop folded into the extended groove, resembling its configuration in the basal state (**Fig. 5D,E**). These changes effectively dismantled the original S1 and S2 sites and significantly weakened the S3 site. Compared to the N-domain groove, the extended groove underwent considerably more pronounced changes (**Fig. 5D,E**).

Taken together, assuming that the mechanistic steps in V2Rpp binding and dissociation are reversible, we propose that the ’p3-in-S5’ and ’p4-in-S5’ states represent intermediate configurations in the V2Rpp binding process. In this mechanism, p3 transiently occupies the partially formed S5 site first, followed by p4, serving as preparatory steps that facilitate the subsequent sliding of p5 into the site, ultimately establishing the strongest interaction between V2Rpp and βarr1 (**Fig. 5F-G**). Once p5 to p8 settle into their designated sites within the N-domain groove (**Fig. 5H**), they allosterically modulate the finger loop to adopt an upward orientation, as described above. The entry of p2 and p3 into the extended groove attracts the middle and “157” loops to their active-state positions, establishing the S2 and S3 sites (**Fig. 5I**). Consequently, the engagement of p1 to p3 with the extended groove likely follows a zippering motion. Together, these sliding and zippering motions correspond to the relatively minor rearrangements in the N-domain groove and the more pronounced changes in the extended groove, respectively.

### The potential functionally relevant conformational ensembles of the βarr1 tail and their anchoring points on the main body

In our TREMD simulations of the V2Rpp-bound active state of βarr1, the dissociation of the βarr1 tail from the N-domain groove allowed it to explore a vast conformational space (**Fig. 6B**). However, analyzing the conformational ensemble and dynamics of the tail was challenging due to the diversity of the sampled conformations. Although we applied both RMSD-based and principal component analysis-based clustering approaches, these methods did not yield tractable results (data not shown) for meaningful insights. The observed highly dynamic nature of the 62-residue βarr1 tail, consistent with the difficulties in experimentally resolving its structure, prompted us to formulate the following hypotheses to guide further analysis: (i) the long tail should not be treated as a single entity but rather as modular, with some segments behaving relatively independently of others; (ii) the interactions with the main body serve as critical anchoring points to facilitate the tail’s functional roles; and (iii) the functionally relevant conformational states of βarr tail may occupy only a small fraction of conformational space sampled in our simulations.

**Figure 6.**
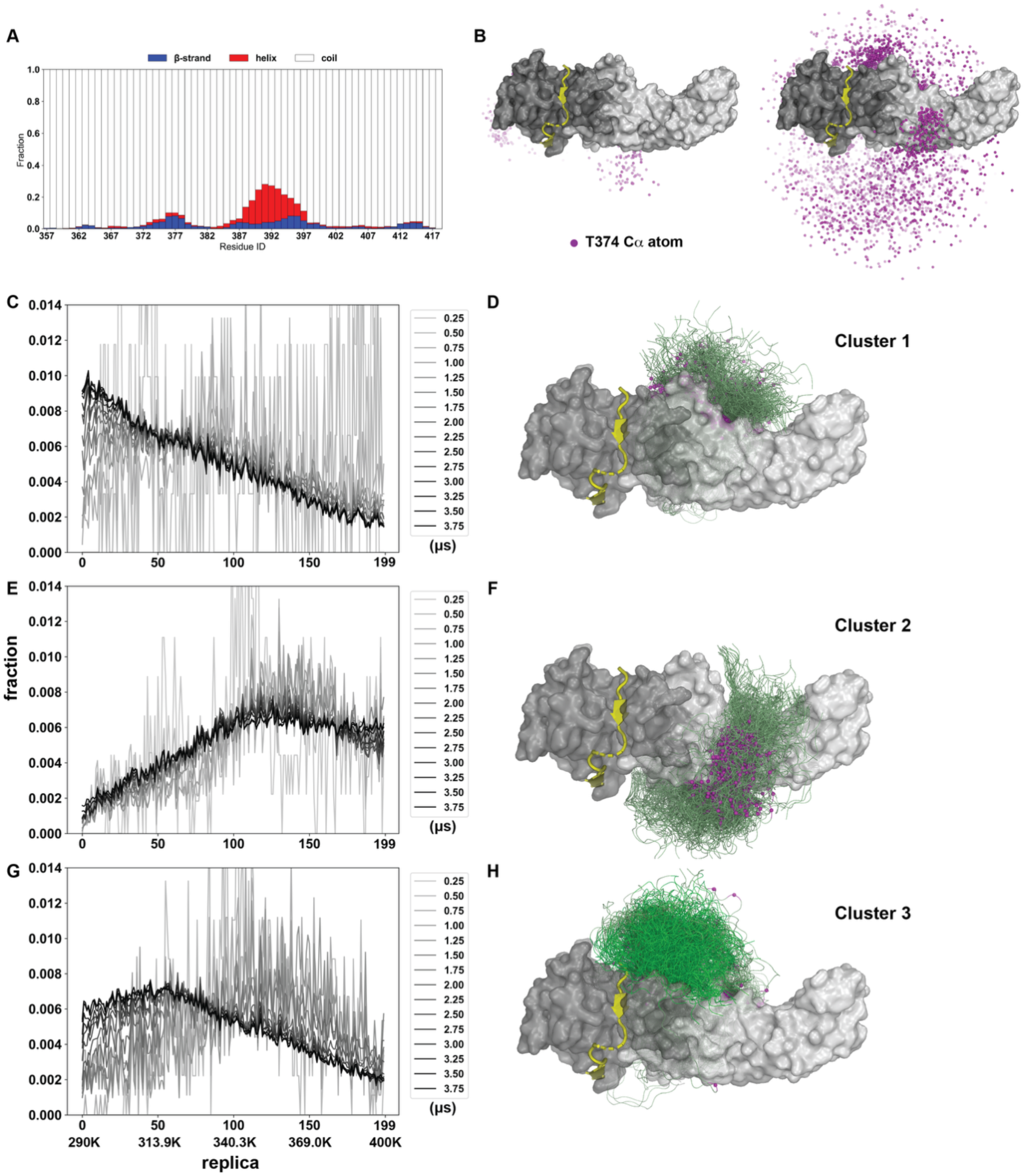
Enhanced sampling of βarr1 C-terminal tail dynamics and identification of metastable substates in V2Rpp-bound TREMD simulations. Per-residue secondary structure analysis of the βarr1 C-terminal tail (residues 357-418) during the V2Rpp-bound βarr1 TREMD simulation reveals transient formation of β-strand and α-helical elements, with the chameleon motif (residues 388-397) frequently adopting helical turns upon release from the N-domain groove. Representative ensembles of the Cα atom of Thr374 from conventional MD and TREMD (panel B) illustrate the substantially broader conformational space accessed by TREMD. Panels C, E, and G show accumulated population distributions of Clusters 1-3 across temperature-ranked replicas, with distributions at selected time points colored from light gray (0.25 µs) to black (3.75 µs). Clusters 1 and 3 are more populated in lower-temperature replicas, supporting their stability and convergence. Panels D, F, and H show representative structural snapshots of Clusters 1-3. The Cα atom of Thr374 is shown as a magenta sphere; the proximal tail segment is colored gray-green, and the chameleon motif is shown in bright green.

Based on these hypotheses, we first analyzed the secondary structure the βarr1 tail during the TREMD simulations and observed transient formation of β-strand and α-helical conformations in specific regions (**Fig. 6A**). Notably, the middle segment of the βarr1 tail, i.e., the chameleon motif, which adopts an extended conformation within the N-domain groove in the basal state, showed a significant tendency to form both β-strand and α-helical conformations when no longer interacting with the groove. This observation suggests that the proximal, middle, and distal segment boundaries, previously defined based on basal state simulations (Asher et al. 2022), can also be applied to analyze βarr1 tail conformations in the active state.

We then calculated all the interactions between the βarr1 tail and the main body across the entire TREMD simulation frame set to identify the residues in both regions with relatively high probabilities of interaction (**Fig. S7**). As expected, several residues in the first half of the proximal segment of the βarr1 tail (residues 357-366), located immediately adjacent to the main body, showed high interaction probability with various main body residues due to their proximity. Moving to more distal parts of the βarr1 tail, we identified two patches of residues frequently interacting with the main body: residues 370-376 in the second half of the proximal segment and residues 392-396 in the chameleon motif. These regions interacted with distinct parts of the main body, namely arrestin switch III (ASwIII, residues 307-316) and the C-loop (residues 242-246), respectively, both in the C domain (**Fig. S7**).

Based on the distances between a selected residue pair from segment 370-376 and AswIII (Thr374 and Arg307, respectively), we identified frames where these residues were in close proximity. These frames were then used as a seed ensemble to gather all similar frames into a cluster (referred to as “Cluster 1”) using an RMSD-based approach (see Methods). We found that this cluster of the βarr1 tail interacted with the side of the main body opposite to the V2Rpp-bound N-domain groove (the ‘back’ side, **Fig. 6D**). An analysis of the number of frames across replicas showed that this cluster was enriched in the low-temperature replicas, indicating it represented an energetically favorable state. Importantly, this distribution trend converged in the later stage (>2.0 µs) of the simulation (**Fig. 6C**). To further validate the energetic preference of “Cluster 1”, we assembled a control cluster using a visually identified residue pair, Asn375 of the proximal segment and Met255 on the “front” side of the C domain. This cluster (“Cluster 2”) represented the interaction of the βarr1 tail with the “front” side of the main body (**Fig. 6F**). In contrast to “Cluster 1”, frames belonging to this cluster tended to have higher distributions in high-temperature replicas, suggesting a high-energy state (**Fig. 6E**). These results suggested that the βarr1 tail was more likely to interact with the “back” side of the main body, consistent with our previous deductions based on the results from a single-molecule FRET study (Asher et al. 2022).

Similarly, for the other pair of frequently interacting patches in the chameleon motif and the C loop, we used a selected residue pair (residues Ala392 and Phe244, respectively) to identify the seed ensemble and clustered similar frames. Similar to “Cluster 1”, this cluster (“Cluster 3”) also showed enrichment in low-temperature replicas, with its distribution trend converging during the later stages of the simulations (**Fig. 6G**). Interestingly, the frames in “Cluster 3” showed that the chameleon motif largely occupied the central crest crevice, which is partly formed by the C loop (**Fig. 6H**). Although the motif was not always in a helical conformation in our simulations (**Fig. 6A**), its occupation of the pocket resembled the binding of the chameleon motif to the central crest crevice observed in the cryo-EM structure of βarr2 (PDB 8J8V) (Maharana et al. 2024). As this crevice serves as the binding site for intracellular loop 2 of GPCRs in the core engagement, its occupation by the chameleon motif is likely to hinder this engagement and to favor tail only engagement.

Thus, our TREMD simulations revealed that the βarr1 tail exhibited a strong tendency to interact with the main body on the “back” side, while the middle segments had a significant probability of binding in the central crest crevice pocket. These interactions are likely to play a key role in modulating βarr1’s interactions with receptors and its functional mechanisms.

## DISCUSSION

The finger loop of arrestins plays a critical role analogous to the C-terminus of Gα proteins in recognizing and interacting with the intracellular crevice of GPCRs, and peptides mimicking either the finger loop or the Gα C-terminus can induce distinct conformational changes in the receptor binding crevice (Szczepek et al. 2014). We found that V2Rpp binding within the N-domain groove allosterically modulated the finger loop, orienting it upward to facilitate the interactions with the intracellular crevice of GPCRs. The critical role of the binding within the N-domain groove but not the extended groove is supported by the observation that phosphopeptides in the V2Rpp-bound βarr2 (PDB 8I10) and the D6Rpp-bound βarr1 and βarr2 (PDBs 8J8Z and 8J8V) structures occupy the N-domain groove but not the extended groove (Maharana et al. 2023, Maharana et al. 2024). Interestingly, in a basal arrestin 1 structure, the C-terminal tail was nearly completely resolved in two of four monomers (PDB 7JSM)(Sander et al. 2022). In these two monomers, the C-terminal tail is at the rim of the extended groove and interacts with the 157 loop, while the extended groove is mainly occupied by the 344 loop of a neighboring monomer. The finger loop in these two monomers adopts an upward orientation, while in the other two monomers (where the C-terminal tail is not proximal to the extended grove), the finger loop folds into the extended groove. These distinct finger loop orientations align with our TREMD simulations, which showed the finger loop orientation depends on occupancy of the extended groove when the N-domain groove is bound to the arrestin C-terminal tail. Together, we found that the binding of phosphopeptides but not the arrestin C-terminal tail in the N-domain groove allosterically modulates the finger loop to prime arrestin for GPCR engagement.

The relative rotation between the N and C domains, quantified by the twist angle, may contribute to the arrestin activation and its interaction with GPCRs. However, no evidence has shown that the activation or receptor engagement requires a specific twist angle as a geometric constraint. For examples, the receptor-arrestin complexes formed without Fab30 exhibited significantly smaller twist angles, such as those of the rhodopsin-arrestin 1 (PDB 5W0P)(Zhou et al. 2017) and the NTSR1-βarr1 (PDB 6UP7)(Huang et al. 2020) complexes, compared to the complexes stabilized by Fab30 (**Tables S2 and S3**). Notably the NTSR1-βarr1 complex bound with Fab30 (PDB 6PWC)(Yin et al. 2019) displays noticeably larger twist angle than its Fab30-free counterpart 6UP7. Similarly, a smaller twist angle was observed for the βarr2 structure activated by the binding of IP6 in the absence of Fab30 (PDB 5TV1)(Chen et al. 2017).

Furthermore, in the absence of Fab30, βarr2 bound with the phosphopeptide derived from the C-terminus of CXCR7, a known activator (PDB 6K3F)(Min et al. 2020), exhibited a twist angle similar to the basal state. Intriguing, the C domain in this structure slightly rotated to the opposite direction, compared to the Fab30-stablized phosphopeptide-bound βarr2 structures (**Tables S2 and S3**). For the arrestin 1 structures that are in the so-called preactivated state, which has the finger loop in an intermediate position between the inactive and active states (Salom et al. 2025), their twist angles resembled the basal state (**Tables S2 and S3**). Thus, it is likely that the finger loop rearrangement occurs relatively independently of inter-domain rotation. Taken together, these findings suggest that the twist angle may be a feature of the Fab30 stabilized state and not arrestin activation per se.

### Author Contribution

VN, JAJ, LS conceptualized the work. VN and LS designed and carried out the computations and developed the analysis approaches. VN, WBA, JAJ, and LS analyzed the data. VN and LS wrote the initial draft, and all authors contributed to finalizing the manuscript.

## Acknowledgements

Support for this research was provided by the National Institute on Drug Abuse–Intramural Research Program, Z1A DA000606 (L.S.), by NIMH R01 MH541397 (J.A.J.) and the Hope for Depression Research Foundation (J.A.J.), and by the Advanced Scientific Computing Research (V.N.), the Office of Science at DOE (No. ERKJZN1). This research used resources of the Oak Ridge Leadership Computing Facility (OLCF) at the Oak Ridge National Laboratory, which is supported by the Office of Science of the U.S. Department of Energy under Contract No. DE-AC05-00OR22725. The TREMD simulations were carried out on the Frontier supercomputer at the OLCF, under the INCITE 2024 Award BIP248. The MD simulations were performed using the NIH high-performance computing Biowulf cluster (https://hpc.nih.gov). Data analysis utilized computational resources on both the Andes cluster at OLCF and Biowulf.

## REFERENCE

Ahn, S., S. K. Shenoy, L. M. Luttrell and R. J. Lefkowitz (2020). "SnapShot: beta-Arrestin Functions." Cell 182(5): 1362–1362 e1361.

Asher, W. B., D. S. Terry, G. G. A. Gregorio, A. W. Kahsai, A. Borgia, B. Xie, A. Modak, Y. Zhu, W. Jang, A. Govindaraju, L. Y. Huang, A. Inoue, N. A. Lambert, V. V. Gurevich, L. Shi, R. J. Lefkowitz, S. C. Blanchard and J. A. Javitch (2022). "GPCR-mediated beta-arrestin activation deconvoluted with single-molecule precision." Cell 185(10): 1661–1675 e1616.

Cahill, T. J., 3rd, A. R. Thomsen, J. T. Tarrasch, B. Plouffe, A. H. Nguyen, F. Yang, L. Y. Huang, A. W. Kahsai, D. L. Bassoni, B. J. Gavino, J. E. Lamerdin, S. Triest, A. K. Shukla, B. Berger, J. t. Little, A. Antar, A. Blanc, C. X. Qu, X. Chen, K. Kawakami, A. Inoue, J. Aoki, J. Steyaert, J. P. Sun, M. Bouvier, G. Skiniotis and R. J. Lefkowitz (2017). "Distinct conformations of GPCR-beta-arrestin complexes mediate desensitization, signaling, and endocytosis." Proc Natl Acad Sci U S A 114(10): 2562–2567.

Chen, Q., N. A. Perry, S. A. Vishnivetskiy, S. Berndt, N. C. Gilbert, Y. Zhuo, P. K. Singh, J. Tholen, M. D. Ohi, E. V. Gurevich, C. A. Brautigam, C. S. Klug, V. V. Gurevich and T. M. Iverson (2017). "Structural basis of arrestin-3 activation and signaling." Nat Commun 8(1): 1427.

Gurevich, V. V. and E. V. Gurevich (2015). "Arrestins: Critical Players in Trafficking of Many GPCRs." Prog Mol Biol Transl Sci 132: 1–14.

Gurevich, V. V. and E. V. Gurevich (2019). "GPCR Signaling Regulation: The Role of GRKs and Arrestins." Front Pharmacol 10: 125.

Gurevich, V. V., E. V. Gurevich and V. N. Uversky (2018). "Arrestins: structural disorder creates rich functionality." Protein Cell 9(12): 986–1003.

Han, M., V. V. Gurevich, S. A. Vishnivetskiy, P. B. Sigler and C. Schubert (2001). "Crystal structure of beta-arrestin at 1.9 A: possible mechanism of receptor binding and membrane Translocation." Structure 9(9): 869–880.

Hastings, W. K. (1970). "Monte Carlo sampling methods using Markov chains and their applications." Biometrika 57(1): 97–109.

Hauser, A. S., M. M. Attwood, M. Rask-Andersen, H. B. Schioth and D. E. Gloriam (2017). "Trends in GPCR drug discovery: new agents, targets and indications." Nat Rev Drug Discov 16(12): 829–842.

He, Q. T., P. Xiao, S. M. Huang, Y. L. Jia, Z. L. Zhu, J. Y. Lin, F. Yang, X. N. Tao, R. J. Zhao, F. Y. Gao, X. G. Niu, K. H. Xiao, J. Wang, C. Jin, J. P. Sun and X. Yu (2021). "Structural studies of phosphorylation-dependent interactions between the V2R receptor and arrestin-2." Nat Commun 12(1): 2396.

Hirsch, J. A., C. Schubert, V. V. Gurevich and P. B. Sigler (1999). "The 2.8 A crystal structure of visual arrestin: a model for arrestin’s regulation." Cell 97(2): 257–269.

Huang, W., M. Masureel, Q. Qu, J. Janetzko, A. Inoue, H. E. Kato, M. J. Robertson, K. C. Nguyen, J. S. Glenn, G. Skiniotis and B. K. Kobilka (2020). "Structure of the neurotensin receptor 1 in complex with beta-arrestin 1." Nature 579(7798): 303–308.

Kahsai, A. W., B. Pani and R. J. Lefkowitz (2018). "GPCR signaling: conformational activation of arrestins." Cell Res 28(8): 783–784.

Kang, Y., X. E. Zhou, X. Gao, Y. He, W. Liu, A. Ishchenko, A. Barty, T. A. White, O. Yefanov, G. W. Han, Q. Xu, P. W. de Waal, J. Ke, M. H. Tan, C. Zhang, A. Moeller, G. M. West, B. D. Pascal, N. Van Eps, L. N. Caro, S. A. Vishnivetskiy, R. J. Lee, K. M. Suino-Powell, X. Gu, K. Pal, J. Ma, X. Zhi, S. Boutet, G. J. Williams, M. Messerschmidt, C. Gati, N. A. Zatsepin, D. Wang, D. James, S. Basu, S. Roy-Chowdhury, C. E. Conrad, J. Coe, H. Liu, S. Lisova, C. Kupitz, I. Grotjohann, R. Fromme, Y. Jiang, M. Tan, H. Yang, J. Li, M. Wang, Z. Zheng, D. Li, N. Howe, Y. Zhao, J. Standfuss, K. Diederichs, Y. Dong, C. S. Potter, B. Carragher, M. Caffrey, H. Jiang, H. N. Chapman, J. C. Spence, P. Fromme, U. Weierstall, O. P. Ernst, V. Katritch, V. V. Gurevich, P. R. Griffin, W. L. Hubbell, R. C. Stevens, V. Cherezov, K. Melcher and H. E. Xu (2015). "Crystal structure of rhodopsin bound to arrestin by femtosecond X-ray laser." Nature 523(7562): 561–567.

Kim, J., S. Ahn, X. R. Ren, E. J. Whalen, E. Reiter, H. Wei and R. J. Lefkowitz (2005). "Functional antagonism of different G protein-coupled receptor kinases for beta-arrestin-mediated angiotensin II receptor signaling." Proc Natl Acad Sci U S A 102(5): 1442–1447.

Kofke, D. A. (2002). "On the acceptance probability of replica-exchange Monte Carlo trials." The Journal of Chemical Physics 117(15): 6911–6914.

Latorraca, N. R., J. K. Wang, B. Bauer, R. J. L. Townshend, S. A. Hollingsworth, J. E. Olivieri, H. E. Xu, M. E. Sommer and R. O. Dror (2018). "Molecular mechanism of GPCR-mediated arrestin activation." Nature 557(7705): 452–456.

Lee, K. H., J. J. Manning, J. Javitch and L. Shi (2023). "A Novel "Activation Switch" Motif Common to All Aminergic Receptors." J Chem Inf Model.

Lee, Y., T. Warne, R. Nehme, S. Pandey, H. Dwivedi-Agnihotri, M. Chaturvedi, P. C. Edwards, J. Garcia-Nafria, A. G. W. Leslie, A. K. Shukla and C. G. Tate (2020). "Molecular basis of beta-arrestin coupling to formoterol-bound beta1-adrenoceptor." Nature 583(7818): 862–866.

Lefkowitz, R. J. (2013). "A brief history of G-protein coupled receptors (Nobel Lecture)." Angew Chem Int Ed Engl 52(25): 6366–6378.

Lefkowitz, R. J. and S. K. Shenoy (2005). "Transduction of receptor signals by beta-arrestins." Science 308(5721): 512–517.

Maharana, J., F. K. Sano, P. Sarma, M. K. Yadav, L. Duan, T. M. Stepniewski, M. Chaturvedi, A. Ranjan, V. Singh, S. Saha, G. Mahajan, M. Chami, W. Shihoya, J. Selent, K. Y. Chung, R. Banerjee, O. Nureki and A. K. Shukla (2024). "Molecular insights into atypical modes of beta-arrestin interaction with seven transmembrane receptors." Science 383(6678): 101–108.

Maharana, J., P. Sarma, M. K. Yadav, S. Saha, V. Singh, S. Saha, M. Chami, R. Banerjee and A. K. Shukla (2023). "Structural snapshots uncover a key phosphorylation motif in GPCRs driving beta-arrestin activation." Mol Cell 83(12): 2091–2107 e2097.

Metropolis, N., A. W. Rosenbluth, M. N. Rosenbluth, A. H. Teller and E. Teller (1953). "Equation of State Calculations by Fast Computing Machines." Journal of Chemical Physics 21(6): 1087–1092.

Michino, M., C. A. Boateng, P. Donthamsetti, H. Yano, O. M. Bakare, A. Bonifazi, M. P. Ellenberger, T. M. Keck, V. Kumar, C. Zhu, R. Verma, J. R. Deschamps, J. A. Javitch, A. H. Newman and L. Shi (2017). "Toward Understanding the Structural Basis of Partial Agonism at the Dopamine D(3) Receptor." J Med Chem 60(2): 580–593.

Min, K., H. J. Yoon, J. Y. Park, M. Baidya, H. Dwivedi-Agnihotri, J. Maharana, M. Chaturvedi, K. Y. Chung, A. K. Shukla and H. H. Lee (2020). "Crystal Structure of beta-Arrestin 2 in Complex with CXCR7 Phosphopeptide." Structure 28(9): 1014–1023 e1014.

Nymeyer, H. (2008). "How Efficient Is Replica Exchange Molecular Dynamics? An Analytic Approach." J Chem Theory Comput 4(4): 626–636.

Perry-Hauser, N. A., J. B. Hopkins, Y. Zhuo, C. Zheng, I. Perez, K. M. Schultz, S. A. Vishnivetskiy, A. I. Kaya, P. Sharma, K. N. Dalby, K. Y. Chung, C. S. Klug, V. V. Gurevich and T. M. Iverson (2022). "The Two Non-Visual Arrestins Engage ERK2 Differently." J Mol Biol 434(7): 167465.

Peterson, Y. K. and L. M. Luttrell (2017). "The Diverse Roles of Arrestin Scaffolds in G Protein-Coupled Receptor Signaling." Pharmacol Rev 69(3): 256–297.

Pierce, K. L., R. T. Premont and R. J. Lefkowitz (2002). "Seven-transmembrane receptors." Nat Rev Mol Cell Biol 3(9): 639–650.

Qi, R., G. Wei, B. Ma and R. Nussinov (2018). "Replica Exchange Molecular Dynamics: A Practical Application Protocol with Solutions to Common Problems and a Peptide Aggregation and Self-Assembly Example." Methods Mol Biol 1777: 101–119.

Rhee, Y. M. and V. S. Pande (2003). "Multiplexed-replica exchange molecular dynamics method for protein folding simulation." Biophys J 84(2 Pt 1): 775-786.

Rosta, E. and G. Hummer (2009). "Error and efficiency of replica exchange molecular dynamics simulations." J Chem Phys 131(16): 165102.

Salom, D., P. D. Kiser and K. Palczewski (2025). "Insights into the Activation and Self-Association of Arrestin-1." Biochemistry 64(2): 364–376.

Sanbonmatsu, K. Y. and A. E. Garcia (2002). "Structure of Met-enkephalin in explicit aqueous solution using replica exchange molecular dynamics." Proteins 46(2): 225–234.

Sander, C. L., J. Luu, K. Kim, D. Furkert, K. Jang, J. Reichenwallner, M. Kang, H. J. Lee, B. T. Eger, H. W. Choe, D. Fiedler, O. P. Ernst, Y. J. Kim, K. Palczewski and P. D. Kiser (2022). "Structural evidence for visual arrestin priming via complexation of phosphoinositols." Structure 30(2): 263–277 e265.

Scheerer, P. and M. E. Sommer (2017). "Structural mechanism of arrestin activation." Curr Opin Struct Biol 45: 160–169.

Shenoy, S. K. and R. J. Lefkowitz (2011). "beta-Arrestin-mediated receptor trafficking and signal transduction." Trends Pharmacol Sci 32(9): 521–533.

Shukla, A. K., A. Manglik, A. C. Kruse, K. Xiao, R. I. Reis, W. C. Tseng, D. P. Staus, D. Hilger, S. Uysal, L. Y. Huang, M. Paduch, P. Tripathi-Shukla, A. Koide, S. Koide, W. I. Weis, A. A. Kossiakoff, B. K. Kobilka and R. J. Lefkowitz (2013). "Structure of active beta-arrestin-1 bound to a G-protein-coupled receptor phosphopeptide." Nature 497(7447): 137–141.

Shukla, A. K., G. H. Westfield, K. Xiao, R. I. Reis, L. Y. Huang, P. Tripathi-Shukla, J. Qian, S. Li, A. Blanc, A. N. Oleskie, A. M. Dosey, M. Su, C. R. Liang, L. L. Gu, J. M. Shan, X. Chen, R. Hanna, M. Choi, X. J. Yao, B. U. Klink, A. W. Kahsai, S. S. Sidhu, S. Koide, P. A. Penczek, A. A. Kossiakoff, V. L. Woods, Jr., B. K. Kobilka, G. Skiniotis and R. J. Lefkowitz (2014). "Visualization of arrestin recruitment by a G-protein-coupled receptor." Nature 512(7513): 218–222.

Smith, J. S., R. J. Lefkowitz and S. Rajagopal (2018). "Biased signalling: from simple switches to allosteric microprocessors." Nat Rev Drug Discov 17(4): 243–260.

Sriram, K. and P. A. Insel (2018). "G Protein-Coupled Receptors as Targets for Approved Drugs: How Many Targets and How Many Drugs?" Mol Pharmacol 93(4): 251–258.

Staus, D. P., H. Hu, M. J. Robertson, A. L. W. Kleinhenz, L. M. Wingler, W. D. Capel, N. R. Latorraca, R. J. Lefkowitz and G. Skiniotis (2020). "Structure of the M2 muscarinic receptor-beta-arrestin complex in a lipid nanodisc." Nature.

Stolzenberg, S., M. Michino, M. V. LeVine, H. Weinstein and L. Shi (2016). "Computational approaches to detect allosteric pathways in transmembrane molecular machines." Biochim Biophys Acta 1858(7 Pt B): 1652–1662.

Sugita, Y. and Y. Okamoto (1999). "Replica-exchange molecular dynamics method for protein folding." Chemical Physics Letters 314(1-2): 141–151.

Szczepek, M., F. Beyriere, K. P. Hofmann, M. Elgeti, R. Kazmin, A. Rose, F. J. Bartl, D. von Stetten, M. Heck, M. E. Sommer, P. W. Hildebrand and P. Scheerer (2014). "Crystal structure of a common GPCR-binding interface for G protein and arrestin." Nat Commun 5: 4801.

Tobin, A. B. (2008). "G-protein-coupled receptor phosphorylation: where, when and by whom." Br J Pharmacol 153 **Suppl 1**: S167–176.

Weis, W. I. and B. K. Kobilka (2018). "The Molecular Basis of G Protein-Coupled Receptor Activation." Annu Rev Biochem 87: 897–919.

Xu, Z. and Z. Shao (2022). "Dynamic mechanism of GPCR-mediated beta-arrestin: a potential therapeutic agent discovery of biased drug." Signal Transduct Target Ther 7(1): 283.

Yin, W., Z. Li, M. Jin, Y. L. Yin, P. W. de Waal, K. Pal, Y. Yin, X. Gao, Y. He, J. Gao, X. Wang, Y. Zhang, H. Zhou, K. Melcher, Y. Jiang, Y. Cong, X. Edward Zhou, X. Yu and H. Eric Xu (2019). "A complex structure of arrestin-2 bound to a G protein-coupled receptor." Cell Res 29(12): 971–983.

Zhou, X. E., Y. He, P. W. de Waal, X. Gao, Y. Kang, N. Van Eps, Y. Yin, K. Pal, D. Goswami, T. A. White, A. Barty, N. R. Latorraca, H. N. Chapman, W. L. Hubbell, R. O. Dror, R. C. Stevens, V. Cherezov, V. V. Gurevich, P. R. Griffin, O. P. Ernst, K. Melcher and H. E. Xu (2017). "Identification of Phosphorylation Codes for Arrestin Recruitment by G Protein-Coupled Receptors." Cell 170(3): 457–469 e413.

