## Supplemental Materials for "Conformational dynamics of the active state of β-arrestin 1"

### METHODS

#### Molecular modeling

The modeling of βarr1 in its basal state was previously described (Asher et al. 2022). The active-state model of βarr1 bound to V2Rpp was constructed following a similar approach. First, homology modeling was performed using Modeller (Webb and Sali 2016), with an active βarr1 structure (PDB: 4JQI) (Shukla et al. 2013) serving as the template. Regions lacking a structural template, including the entire C-terminal tail, were modeled ab initio using Modeller (Webb and Sali 2016) and further refined with ROSETTA (Stein and Kortemme 2013) (see (Asher et al. 2022)). The terminal residues of the generated models were assigned physiologically relevant charge states, with the first residue positively charged and the last negatively charged, without capping. The prepared structures were solvated in a simulation water box using CHARMM-GUI (Jo et al. 2008), employing the TIP3P water model (Jorgensen et al. 1983, Neria et al. 1996). Sodium ($Na^+$) and chloride ($Cl^-$) ions were added to neutralize the system and achieve a physiological salt concentration of 0.15 M. To accurately sample the conformational ensemble of the βarr1 C-terminal tail, which is expected to undergo unwinding and rewinding transitions during TREMD, a sufficiently large simulation box ($143 \times 143 \times 143$ Å$^3$) was used. This ensured that the C-tail could fully extend without interacting with its periodic images. The total system size was approximately 287,000 atoms.

#### Molecular dynamics simulations

MD simulations of the βarr1 systems were carried out with NAMD 3.0b7 (Phillips et al. 2005, Phillips et al. 2020) using the CHARMM36m force field (Huang et al. 2017). The simulation systems were minimized and equilibrated with restraints on the heavy atoms of the protein backbone. For both the equilibrations and the following MD and TREMD production runs, the constant temperature 310 K was maintained by Langevin dynamics, 1 atm constant pressure was achieved with the Langevin piston method (Feller et al. 1995). A switching distance of 10.0 Å was applied to gradually reduce the strength of the non-bonded interactions to 0 at a cutoff distance of 12 Å. The electrostatics interactions were computed using the particle-mesh Ewald summation method (Essmann et al. 1995). An integration timestep of 2.0 fs was used for equilibration stage. The isothermal-isobaric (NPT) ensemble was used in a periodic boundary condition in equilibration. The production MD and TREMD runs were carried out in isothermal-isochoric (NVT) ensemble with the Langevin dynamics with the damping constant of 1 $ps^{-1}$, without any restraints. To use 4 fs timestep in production runs, we apply hydrogen mass repartitioning (see text) (Hopkins et al. 2015).

To properly simulate the interactions between the phosphate and the $Na^+$ ion in the water phase, we applied the NBFIX correction of Yoo and Aksimentiev (Yoo and Aksimentiev 2012).

To accelerate the simulations, we implemented the hydrogen mass repartitioning (HMR) technique, enabling the use of a longer timestep of 4 femtoseconds (fs) instead of 2 fs used without HMR (see Methods) (Hopkins et al. 2015). To assess the stability and reliability of this technique in the simulations, we collected both MD and TREMD simulation trajectories using HMR (**Table S1**) and compared the RMSD and twist angle distributions of these trajectories to those from simulations without HMR (referred to as non-HMR). Our results demonstrated that, although the HMR and non-HMR MD simulations appeared to slightly diverge in individual trajectories to explore different conformational space, the HMR technique had negligible impact on the simulated dynamics in TREMD (**Fig. S6**). Therefore, the divergence observed in MD was likely due to inadequate sampling rather than the impact of HMR. We then proceeded with the TREMD simulations of the active state using HMR and averaged the results of HMR and non-HMR simulations of the basal state for comparisons between these two states.

**Temperature replica-exchange molecular dynamics (TREMD) simulations**

For TREMD simulations, we chose a temperature ladder following the equation,

$$T_i = T_{min}\left(\frac{T_{max}}{T_{min}}\right)^{\frac{i}{N-1}},$$

where $T_{min}$ = 290 K, $T_{max}$ = 400 K, *i* is the replica index starting (ranging from 0 to N-1), and N = 200 is the total number of replicas. The upper temperature ($T_{max}$ = 400 K) was selected to enable adequate *cis-trans* isomerization of peptide bonds (Neale et al. 2016), while preventing the bound V2Rpp from undergoing full dissociation from the main body. The replica exchange acceptance probability in TREMD sampling depends on temperature spacing between adjacent replicas (Kofke 2002). To maximize sampling efficiency, we employed 200 replicas across the temperature range ($T_{max}$ – $T_{min}$), achieving an average exchange rate of 41%. This combination of high exchange rate and sufficiently elevated $T_{max}$ accelerates sampling convergence by promoting rapid exploration of conformational space (Zhang et al. 2005, Periole and Mark 2007, Rosta and Hummer 2009).

Exchange attempts were performed every 1000 steps. Frames were saved every 0.5 ns. The force fields and other simulation configurations were the same as the MD simulations (see above). The lengths of the simulations for each simulated condition were summarized in **Table S1**.

**Residue ranges used in the conformational analysis**

Because the RMSD and twist angle analysis are sensitive to the residue sets chosen for the superposition and/or the rotational angle calculations (data not shown), we used only C$\alpha$ atoms from the most stable regions in the N and C domains. To identify these stable regions, we developed an iterative root-mean-square fluctuation (RMSF) calculation procedure. For the N domain, for instances, we initially superimposed the residues of the entire N domain for all the frames a trajectory, selected the top 70 residues with the lowest RMSF for the subsequent round of superposition, and repeat this process until the composition of the top 70 residues stabilized. We then determined the overlapping stable residues across three basal-state and three V2Rpp-bound-state TREMD trajectories, resulting in 63 residues that constitute the stable regions of the N domain. The same approach applied to the C domain identified 66 residues forming its stable regions.

Specially for bovine βarr1, these residues are 8 to 12, 18 to 26, 36 to 43, 53 to 61, 79 to 82, 108, 111 to 120, 123 to 124, 142 to 149, and 163 to 169 in the N domain, and residues 198 to 203, 207 to 210, 213 to 221, 231 to 239, 252 to 256, 268 to 274, 288 to 289, 300 to 303, 318 to 327, 343 to 352 in the C domain. The aligned βarr1 residues in other species, as well as βarr2 and arrestin 1, were identified from the sequence alignment shown in **Fig. S?** for the twist angle calculations shown in **Table S3**.

Note that these residues are predominantly from the structured regions of the main body but do not encompass all the structured regions, some parts of which were flexible during the simulations. For dividing the entire main body, including both the structured and loop regions, for the PIA analysis, we defined subsegments by integrating those previously described (Hirsch et al. 1999, Gurevich et al. 2018), adjusting them based on visual inspection of experimentally determined structures, and subdividing long β-strands into shorter subsegments to enhance resolution. Detailed definitions of these subsegments are provided in **Table S5**.

**Definition of the twist angle between the N and C domains**

Our twist angle definition was partially inspired by that reported by Latorraca and Dror (Latorraca et al. 2018), and was facilitated with additional details and code provided through personal communication with the authors. We first aligned two given two arrestin structures (or models), one designated as the reference and the other as the target, using the selected C$\alpha$ atoms from their N domains (see above). A rotation angle was then derived from the selected

Cα atoms of their C domains. Specifically, in a rotational space, we determined a transformation matrix *R* that minimizes

$$L = \sum_i |\vec{r}_i' - R\vec{r}_i|^2,$$

where $\vec{r}_i$ and $\vec{r}_i'$ are 3D coordinates of each selected C domain atoms in the target and reference structures, respectively. *R* can be presented by means of a quaternion (*q*), which was defined as follows,

$$q(\gamma, \boldsymbol{n}) = \{q_0, q_1, q_2, q_3\} = q_0 + q_1\boldsymbol{x} + q_2\boldsymbol{y} + q_3\boldsymbol{z} = \{\cos\left(\frac{\gamma}{2}\right), \sin\left(\frac{\gamma}{2}\right)\boldsymbol{n}\},$$

where the unit vector ***n*** is the rotational axis and $\gamma$ is the rotational angle. Then *R* is expressed in *q* as

$$R = \{R_{ij}\} = \begin{bmatrix} R_{00} & R_{01} & R_{02} \\ R_{10} & R_{11} & R_{12} \\ R_{20} & R_{21} & R_{22} \end{bmatrix} = \begin{vmatrix} q_0^2 + q_1^2 - q_2^2 - q_3^2 & 2(q_1q_2 - q_0q_3) & 2(q_1q_3 + q_0q_2) \\ 2(q_1q_2 + q_0q_3) & q_0^2 - q_1^2 + q_2^2 - q_3^2 & 2(q_2q_3 - q_0q_1) \\ 2(q_1q_3 - \text{▯▯}_0q_2) & 2(q_2q_3 + q_0q_1) & q_0^2 - q_1^2 - q_2^2 + q_3^2 \end{vmatrix}$$

$$= \begin{vmatrix} 1 - 2(q_2^2 + q_3^2) & 2(q_1q_2 - q_0q_3) & 2(q_1q_3 + q_0q_2) \\ 2(q_1q_2 + q_0q_3) & 1 - 2(q_1^2 + q_3^2) & 2(q_2q_3 - q_0q_1) \\ 2(q_1q_3 - q_0q_2) & 2(q_2q_3 + q_0q_1) & q_0^2 - 2(q_1^2 + q_2^2) \end{vmatrix}$$

Conversely, the quaternion components were recovered as follows:

$$q_0 = \frac{1}{2}\sqrt{1 + R_{00} + R_{11} + R_{22}} = \cos\left(\frac{\gamma}{2}\right)$$

$$q_1 = \frac{1}{2}\sqrt{1 + R_{00} - R_{11} - R_{22}} \times \mathrm{sign}(R_{21} - R_{12}) = x\sin\left(\frac{\gamma}{2}\right)$$

$$q_2 = \frac{1}{2}\sqrt{1 - R_{00} + R_{11} - R_{22}} \times \mathrm{sign}(R_{02} - R_{20}) = y\sin\left(\frac{\gamma}{2}\right)$$

$$q_3 = \frac{1}{2}\sqrt{1 - R_{00} - R_{11} + R_{22}} \times \mathrm{sign}(R_{10} - R_{01}) = z\sin\left(\frac{\gamma}{2}\right)$$

Therefore, we can compute the rotational angle $\gamma$ and the unit vector ***n*** as

$$\gamma = 2\cos^{-1}(q_0)$$

$$\boldsymbol{n} = \frac{1}{\sin\left(\frac{\gamma}{2}\right)}\{q_1, q_2, q_3\}$$

Because $\gamma$ itself does not define the direction of rotation, the twist angle was introduced as the projection of the rotation angle onto the plane perpendicular to the vector connecting the center of mass (COM) of the N domain to the COM of the C domain in the reference structure (**Fig. S5**). Thus, a positive twist angle indicates a counter-clockwise rotation of the target structure's C domain relative to the reference when viewed from the COM of the N domain toward that of C domain.

In calculating the twist angles for the MD or TREMD simulations of βarr1, we used either the basal-state structure 1ZSH or the active-state structure 4JQI as the references (see text).

**Clustering of the βarr1 tail in the TREMD results of V2Rpp-bound state**

To identify functionally relevant conformational ensembles of the βarr1 tail, we first constructed a contact frequency matrix between βarr1 tail and main body residues. Specifically, residue-residue contacts were evaluated between residues 357 to 418 of the tail and residues 1 to 356 of the main body. For each pairwise combination, a normalized contact ratio was calculated, representing the frequency of contact formation over the simulation time. This yielded a residue-residue ratio matrix that reflects the spatial distribution and temporal persistence of tail-main body interactions.

To extract persistent interaction patterns, a maximum filter with a footprint size of 5×5 was applied to the ratio matrix to identify local maxima. Maxima with contact ratios above a heuristic threshold of 0.1 were retained, ensuring that selected interaction regions were both spatially localized and statistically enriched (Fig. S7). These maxima defined specific residue pairs used for subsequent frame extraction.

For each selected residue pair, trajectory frames were identified in which the inter-residue distance was below a defined cutoff of 15 Å. These frames were collected across all replicas and binned over time to produce block-wise interaction statistics. Blocks showing temporal enrichment of these contacts were considered to reflect stable or emerging interaction modes.

Representative frames from highly enriched blocks were used as seed conformations to initiate clustering. RMSD-based clustering was then performed on conformations surrounding each seed, using the heavy atoms of a localized segment of the βarr1 tail. These segments, typically spanning 15 to 16 residues, were selected to capture the structural context of the anchoring interaction while excluding unrelated tail flexibility. This targeted approach improved sensitivity to conformational differences relevant to specific anchoring modes. Clusters were evaluated based on their distribution across temperature replicas and their accumulation over simulation time (Fig. 6). Clusters that were enriched in low-temperature replicas and showed increasing population in the later stages of the TREMD simulations were considered to represent low-energy, functionally relevant conformational ensembles. Control clusters involving alternate anchoring patterns were also analyzed to compare their energetic preference and structural stability.

### SUPPLEMENTAL RESULTS

#### Benchmarking TREMD for optimal exchange efficiency and enhanced sampling

To determine the necessary number of replicas required to obtain an adequate exchange acceptance ratio among replicas so to achieve optimal sampling speed-up and convergence, we first performed two benchmarking TREMD simulations, each lasting 80 ns. One simulation utilized 100 replicas, while the other employed 200 replicas on the simulation system. In these simulations, we selected a lower-bound temperature of 290K and an upper-bound temperature of 400K for our simulations, a choice inspired by the work of Sanbonmatsu and Garcia (Sanbonmatsu and Garcia 2002). This temperature selection serves two purposes: firstly, to reduce the occurrence of completely unfolding events in the entire complex, which are not functionally relevant (Dobson et al. 1998, Karplus 2011); secondly, to enhance transitions among relevant structures among replicas, facilitating the exploration of the desired conformational space. The analysis of these two simulations showed approximately uniform acceptance ratio across all replicas within a simulation (10 ± 2% and 41 ± 2% for the 100-replica and 200-replica simulations, respectively). The 200-replica simulation yields a better propagation in the temperature space as demonstrated in **Fig. S1**, in which during 80 ns the configurations 17 unsorted replicas in the 200-replica simulation are found to have visited the entire temperature ladder of 290-400 K, while only three replicas in the 100-replica simulation have achieved that, indicating a significantly more extensive sampling in the 200-replica run.

To further characterize whether the 200-replica TREMD simulation yields better sampling than the 100-replica run even within 80 ns, we examined the frequency and extent and a functionally relevant rare event, i.e., the partial dissociation of the V2Rpp from the extended groove. We found that 38 out of the 50 replicas in the lowest temperatures (290-314.4K, after resorting) have P1-P3 of V2Rpp dissociated but only after 70 ns (**Fig. S2**), while this dissociation did not happen in the 100-replica run. Thus, we chose the 200-replica system for the production simulations.

**Estimation of the enhanced sampling by TREMD**

TREMD simulations leverage the Metropolis-Hastings algorithm to sample rare events that are more accessible at higher temperatures. Thus, certain rare events of interests can be observed to occur with significantly higher frequency in TREMD compared to classical MD simulations, demonstrating an enhanced sampling of these rare events. To quantify the enhancement in sampling efficiency achieved by TREMD, we compared occurrences of rare-event transitions in TREMD and MD trajectories at the same temperature 310K. Identifying specific rare events that represent the slowest timescales of a system's dynamics is often challenging. A common strategy involves finding a space or a set of observables that can be effectively represented or approximated by Markovian processes (Noe and Fischer 2008). In Markovian transitions between two states, A and B, where B is the rarer state with much shorter lifetime than A, the numbers of transitions in a MD trajectory of length $\tau_{MD}$ are given by $n_{A\text{->}B}$ and $n_{B\text{->}A}$. For MD trajectories that do not exhibit Markovian behavior, Markovian state modeling can approximate Markovian dynamics using an appropriate lag time $\tau_{lag}$. A key quantity characterizing Markovian dynamics is the unique timescale $\tau$, which corresponds to the slowest mode with the first eigenvalue, $e^{-\tau_{lag}/\tau}$, of the Markovian transition matrix (Ngo et al. 2023). The timescale of the two-state system is given by

$$\tau_{\mathrm{sMD}} = -\frac{dt}{\log(1-p_{A\to B}-p_{B\to A})}$$ (Ngo et al. 2023),

where $dt$ is a timestep, and the transition probabilities are defined as

$$p_{A\to B} = dt \times n_{A\to B}/\tau_{\mathrm{MD}},$$

$$p_{B\to A} = dt \times n_{B\to A}/\tau_{\mathrm{MD}},$$

where $n_{A\to B}$ and $n_{B\to A}$ are numbers of transitions between states A and B. For rare events, where $p_{A\to B}$and $p_{B\to A}$ are often small, the timescale simplifies to

$$\tau_{\mathrm{sMD}} \sim dt/(p_{A\to B} + p_{B\to A}) = \frac{\tau_{\mathrm{MD}}}{n_{\mathrm{A\to B}} + n_{\mathrm{B\to A}}}.$$

For a TREMD trajectory of length $\tau_{\mathrm{TREMD}}$, the corresponding timescale is

$$\tau_{\mathrm{sTREMD}} \sim \frac{\tau_{\mathrm{TREMD}}}{N_{\mathrm{A\to B}} + N_{\mathrm{B\to A}}},$$

where $N_{\mathrm{A\to B}} + N_{\mathrm{B\to A}}$ is the total number of the transitions between the two states observed in TREMD. The sampling speed-up is then defined as

$$\alpha_{A\leftrightarrow B} = \frac{\tau_{\mathrm{sMD}}}{\tau_{\mathrm{sTREMD}}} \cong \frac{\tau_{\mathrm{MD}}}{\tau_{\mathrm{TREMD}}} \times \frac{N_{\mathrm{A\to B}} + N_{\mathrm{B\to A}}}{n_{\mathrm{A\to B}} + n_{\mathrm{B\to A}}}.$$

If a TREMD setup, such as one with a small number of replicas over a large temperature range, does not yield more transitions than MD of similar simulation lengths, the ratio $\frac{N_{\mathrm{A\to B}} + N_{\mathrm{B\to A}}}{n_{\mathrm{A\to B}} + n_{\mathrm{B\to A}}}$ would approach 1. For an MD and a TREMD trajectories with similar lengths $\tau_{\mathrm{MD}} \sim \tau_{\mathrm{TREMD}}$, $\alpha_{A\leftrightarrow B}$ depends on the frequencies of the transitions in both MD and TREMD, which is influenced by the temperature ladder and potential energy landscape.

The value of $\alpha_{A\leftrightarrow B}$ can strongly depend on the choice of the states A and B especially when working with limited data sets. In some cases, transitions between states A and B may be sampled in TREMD but not in short MD trajectories, rendering $\alpha_{A\leftrightarrow B}$ undefined. Additionally, assessing the convergence of transition counts in both MD and TREMD remains challenging given computational resource constraints. To estimate the order of magnitude of the speed-up, we identify the state pairs (A, B) that yield the largest speed-up, based on the available MD and TREMD data. For events where no rare transitions are observed in MD (i.e., $n_{A\to B}$ = $n_{B\to A}$ = 0), we calculate the speed-up as $\frac{\tau_{\mathrm{MD}}}{\tau_{\mathrm{TREMD}}}(N_{\mathrm{A\to B}} + N_{\mathrm{B\to A}})$.

We observed that the helical content (HC, defined as the number of residues in helical conformations) of finger-loop and the chameleon motif follows Markovian behavior. To define the Markovian states, we designated state A as having HC below a threshold $HC_a$ and state B as having HC ≥ $HC_a$. By varying $HC_a$, we generated multiple sets of Markovian states A and B to find the largest speed-up based on the aforementioned assumption. In our MD trajectories, and also previously reported MD simulations (Latorraca et al. 2020), the finger loop could briefly adopt a helical conformation with a maximum HC of 4 over ~4-5 μs. In contrast, the TREMD simulations efficiently sampled the finger loop, capturing numerous transitions between HC ≤ 4 and HC > 4 (see **Fig. 5A**). Given these observations, we defined state A for the finger loop as HC ≤ 4 and state B as HC > 4. Consequently, the estimated sampling speed-up of TREMD compared to two independent MD trajectories totaling 7.8 μs was approximately 73-fold. For the chameleon motif, the MD simulations sampled its helical conformation with a maximum HC of 9. We found that TREMD achieved an estimated sampling speed-up of approximately 63-fold for the transition between HC ≤ 9 and HC > 9. Overall, TREMD accelerated the sampling of helical conformations of these loops by roughly two orders of magnitude compared to conventional MD simulations. Based on these findings, we infer that TREMD enhances the sampling of the overall dynamics of βarr1 by a similar factor.

#### Development and evaluation of a twist angle metric for arrestin activation

We aimed to develop an angular metric to generally evaluate the rotational rigid-body movement between the N and C domains during the transition of arrestins from the basal to the active state. Ideally, this metric should have a sufficient dynamic range to capture various extents of movement, provide directional information, and allow comparisons across different arrestin subtypes. To assess the performance of the twist angle we defined from these perspectives, we calculated the twist angle between pairs of 32 experimentally determined arrestin structures, including representative examples of arrestin 1, βarr1, and βarr2 in various states (**Table S2**).

For any given pair of arrestin structures, structure1 and structure2, the twist angles calculated for “structure1 – structure2” and “structure2 – structure1” exhibit similar magnitudes but opposite signs, indicating that the metric accurately reflects the direction of rotation (**Table S3**). The approximately 20° twist angles between the basal and active states are comparable to previously reported values (Shukla et al. 2013, Latorraca et al. 2018, Maharana et al. 2024). Notably, the IP6-bound βarr2 structure (PDB 5TV1), which is in a receptor-independent activated state (Chen et al. 2017), displays a moderate twist angle relative to the basal state, smaller than those observed in phosphopeptide-bound arrestins. This highlights the sensitivity of our twist angle metric in distinguishing activation mechanisms. Furthermore, the relatively small twist angle values observed for “basal – basal” and “active – active” comparisons, along with similar extents of differences across various arrestin subtypes, demonstrate that our twist angle effectively captures the conformational changes common to arrestins (**Table S3**).

Notably, although the non-projected rotational angles (see Methods) generally exhibit larger absolute values (**Table S4**), the differences in twist angles between certain βarr1 and βarr2 pairs are even greater than those observed for the corresponding rotational angles, such as those associated with the structure 6K3F. This suggests that the rotational angles may include components that are not necessarily critical.

### SUPPLEMENTAL TABLES

#### Table S1. Summary of MD and TREMD simulations.

Note that each REMD simulation was carried out using 200 replicas, employing the 290 – 400 K temperature ladder described in the Methods section.

| state | MD | | REMD | |
|---|---|---|---|---|
| | HMR | length (ns) | HMR | Length/replica (ns) |
| basal | - | 2000 | - | 1111 |
| | + | 2000 x2 | + | 1122 |
| v2rpp-bound | - | 4000 x2 | | |
| | + | 4000 | + | 3800 |

**Table S2. Experimentally determined arrestin structures included in the twist angle analysis.**

Synonyms, acronyms, and abbreviations used are as follows: subtype — visual arrestin (arr1), β-arrestin 1 (arr2), β-arrestin 2 (arr3); state — active (a), basal (b), complexed with receptor (c), bound with IP6 (i), bound only with phosphopeptide (p), preactivated (pa); receptor — serotonin 2B receptor (5HT2B), chemokine receptor CXCR7 (CXCR7), β1 adrenergic receptor (ADRB1), cannabinoid receptor 1 (CB1R), muscarinic M2 receptor (ACM2), neurotensin receptor 1 (NTSR1), rhodopsin (RHO), vasopressin V2 receptor (V2R).

| PDB | resolution (Å) | released year | subtype | species | chain | state | receptor\| tail | receptor\| core | finger loop | T4L or antibody | reference |
|---|---|---|---|---|---|---|---|---|---|---|---|
| 3UGU | 1.85 | 2012 | arr1 | bovine | A | pa | | | missing | N/A | (Granzin et al. 2012) |
| 4ZRG | 2.70 | 2015 | arr1 | bovine | A | pa | | | missing | N/A | (Granzin et al. 2015) |
| 9C6E | 1.40 | 2024 | arr1 | bovine | A | pa | | | disordered | N/A | (Salom et al. 2025) |
| 7JSM | 2.50 | 2021 | arr1 | bovine | A | b | | | disordered | N/A | (Sander et al. 2022) |
| 7JTB | 2.60 | 2021 | arr1 | bovine | A | b\|i | | | disordered | N/A | (Sander et al. 2022) |
| 1CF1 | 2.80 | 1999 | arr1 | bovine | A | b | | | disordered | N/A | (Hirsch et al. 1999) |
| 4J2Q | 3.00 | 2013 | arr1 | bovine | A | a | | | disordered | N/A | (Kim et al. 2013) |

| | | | | | | | | | | | |
|---|---|---|---|---|---|---|---|---|---|---|---|
| 5W0P | 3.01 | 2017 | arr1 | mouse | A | c | RHO | RHO | helical | T4L | (Zhou et al. 2017) |
| 5DGY | 7.70 | 2016 | arr1 | mouse | A | c | | RHO | helical | T4L | (Zhou et al. 2016) |
| 1ZSH | 2.90 | 2006 | arr2 | bovine | A | b\|i | | | disordered | N/A | (Milano et al. 2006) |
| 1G4M | 1.90 | 2001 | arr2 | bovine | A | b | | | disordered | N/A | (Han et al. 2001) |
| 1JSY | 2.90 | 2002 | arr2 | bovine | A | b | | | disordered | N/A | (Milano et al. 2002) |
| 2WTR | 2.90 | 2009 | arr2 | bovine | A | b | | | disordered | N/A | |
| 4JQI | 2.60 | 2013 | arr2 | rat | A | p | V2R | | disordered | Fab30 | (Shukla et al. 2013) |
| 8J8Z | 3.40 | 2023 | arr2 | rat | A | p | D6R | | helical | Fab30 | (Maharana et al. 2024) |
| 8JA3 | 3.94 | 2023 | arr2 | rat | A | p | C3aR | | missing | Fab30 | (Maharana et al. 2024) |
| 8JAF | 3.10 | 2023 | arr2 | bovine | A | p | ACM2 | | missing | Fab30 | (Maharana et al. 2024) |
| 6TKO | 3.30 | 2020 | arr2 | human | B | c | V2R | ADRB1 | disordered | Fab30 | (Lee et al. 2020) |
| 6U1N | 4.00 | 2020 | arr2 | rat | C | c | V2R | ACM2 | disordered | Fab30 | (Staus et al. 2020) |

| | | | | | | | | | | | |
|---|---|---|---|---|---|---|---|---|---|---|---|
| 6UP7 | 4.20 | 2020 | arr2 | human | B | c | | NTSR1 | helical | N/A | (Huang et al. 2020) |
| 6PWC | 4.90 | 2019 | arr2 | human | A | c | | NTSR1 | helical | Fab30 | (Yin et al. 2019) |
| 7R0C | 4.73 | 2022 | arr2 | human | C | c | V2R | V2R | twisted β-hairpin | Fab30 | (Bous et al. 2022) |
| 7SRS | 3.30 | 2022 | arr2 | human_1B | C | c | | 5HT2B | helical | Fab30 | (Cao et al. 2022) |
| 8WRZ | 3.60 | 2024 | arr2 | human | A | c | V2R | CB1R | helical | Fab30 | (Wang et al. 2024) |
| 8WU1 | 3.20 | 2024 | arr2 | bovine | C | c | V2R | CB1R | disordered | Fab30 | (Liao et al. 2023) |
| 6K3F | 2.30 | 2020 | arr3 | rat | A | p | CXCR7 | | helical | N/A | (Min et al. 2020) |
| 8J9K | 3.50 | 2023 | arr3 | rat | C | b | | | disordered | Fab6 | (Maharana et al. 2024) |
| 3P2D | 3.00 | 2011 | arr3 | bovine | A | b | | | disordered | N/A | (Zhan et al. 2011) |
| 5TV1 | 2.40 | 2017 | arr3 | bovine | A | a\|i | | | helical | N/A | (Chen et al. 2017) |
| 8I10 | 3.96 | 2023 | arr3 | bovine | A | p | V2R | | disordered | Fab30 | (Maharana et al. 2023) |
| 8J8R | 2.90 | 2023 | arr3 | bovine | A | p | ACM2 | | disordered | Fab30 | (Maharana et al. 2024) |

| | | | | | | | | | | | |
|---|---|---|---|---|---|---|---|---|---|---|---|
| 8J8V | 3.22 | 2023 | arr3 | bovine | A | p | D6R | | disordered | Fab30 | (Maharana et al. 2024) |

**Table S3. Twist angles among experimentally determined arrestin structures.**

Only the structures with resolution of 5.0Å or better listed in Table S2 were included in the twist angle analysis. Structure 8JA3 was excluded due to an incomplete N domain. The structures listed vertically in the Table serve as the reference structures for each calculation. Depending on the reference structure chosen, twist angles calculated between a pair of structures have opposite signs and similar (but not identical) magnitudes, as the plane onto which the rotational angle is projected to derive the twist angle depends on the reference structure (see Methods). For example, using 1ZSH as a reference, the angle for 4JQI – 1ZSH is 16.9°, while using 4JQI as a reference, the angle for 1ZSH – 4JQI is -14.3°.

Structure 4J2Q is in an active conformation despite lacking a bound phosphopeptide or a receptor. The C7pp-bound βarr2 structure 6K3F exhibited a unique C domain conformation distinct from both the basal and Fab30-stablized conformations. It shows twist angles to other structures like a basal-state structure and is, in fact, rotated even further than the basal-state structures relative to the Fab30-stabilized active structures.

| | 3UGU | 4ZRG | 9C6E | 7JSM | 7JTB | 1CF1 | 4J2Q | 5W0P | 1ZSH | 1G4M | 1JSY | 2WTR | 4JQI | 8J8Z | 8JAF | 6TKO | 6U1N | 6UP7 | 6PWC | 7R0C | 7SRS | 8WRZ | 8WU1 | 6K3F | 8J9K | 3P2D | 5TV1 | 8I10 | 8J8R | 8J8V |
|---|---|---|---|---|---|---|---|---|---|---|---|---|---|---|---|---|---|---|---|---|---|---|---|---|---|---|---|---|---|---|
| 3UGU | 0.0 | 0.4 | 1.7 | 1.3 | 1.4 | 0.9 | 15.4 | 15.2 | 5.4 | 4.3 | 5.4 | 4.2 | 21.8 | 21.2 | 20.0 | 19.8 | 19.9 | 17.2 | 19.3 | 20.4 | 20.3 | 20.2 | 18.8 | -9.3 | 3.0 | 0.8 | 16.8 | 20.1 | 19.0 | 18.4 |
| 4ZRG | -0.4 | 0.0 | 1.3 | 0.9 | 1.0 | 0.5 | 14.9 | 14.8 | 4.9 | 3.8 | 5.0 | 3.7 | 21.4 | 20.8 | 19.6 | 19.4 | 19.5 | 16.8 | 18.9 | 20.0 | 19.9 | 19.8 | 18.4 | -9.7 | 2.5 | 0.3 | 16.4 | 19.7 | 18.6 | 18.0 |
| 9C6E | -1.7 | -1.3 | 0.0 | -0.4 | -0.2 | -0.7 | 13.6 | 13.5 | 3.7 | 2.6 | 3.8 | 2.5 | 20.0 | 19.5 | 18.2 | 18.0 | 18.2 | 15.4 | 17.6 | 18.6 | 18.6 | 18.4 | 17.0 | -11.0 | 1.3 | -0.8 | 15.1 | 18.3 | 17.2 | 16.7 |
| 7JSM | -1.3 | -0.9 | 0.4 | 0.0 | 0.2 | -0.2 | 15.1 | 14.2 | 4.2 | 2.9 | 4.2 | 2.7 | 20.3 | 20.0 | 18.9 | 18.4 | 18.5 | 15.0 | 17.9 | 18.7 | 18.8 | 18.6 | 17.4 | -10.9 | 2.0 | 0.0 | 14.4 | 18.8 | 17.4 | 17.1 |
| 7JTB | -1.5 | -1.1 | 0.2 | -0.2 | 0.0 | -0.4 | 15.0 | 14.0 | 4.0 | 2.7 | 4.1 | 2.5 | 20.2 | 19.8 | 18.7 | 18.2 | 18.3 | 14.8 | 17.7 | 18.5 | 18.6 | 18.4 | 17.2 | -11.2 | 1.8 | -0.3 | 14.2 | 18.6 | 17.2 | 16.9 |
| 1CF1 | -1.4 | -1.0 | 0.4 | 0.2 | 0.4 | 0.0 | 14.2 | 14.3 | 4.5 | 3.1 | 4.6 | 2.9 | 21.0 | 20.2 | 19.0 | 19.1 | 19.2 | 16.2 | 18.5 | 19.6 | 19.6 | 19.4 | 18.0 | -10.4 | 2.1 | 0.0 | 15.5 | 19.3 | 18.3 | 17.5 |
| 4J2Q | -15.3 | -14.9 | -13.5 | -13.3 | -12.9 | -13.3 | 0.0 | -0.5 | -8.4 | -10.4 | -8.3 | -10.9 | 5.7 | 5.3 | 4.2 | 3.7 | 3.8 | 0.6 | 3.2 | 4.2 | 4.2 | 4.0 | 2.7 | -24.6 | -11.0 | -13.0 | 0.0 | 4.1 | 2.8 | 2.5 |
| 5W0P | -15.3 | -14.8 | -13.5 | -13.2 | -12.9 | -13.1 | 0.4 | 0.0 | -8.9 | -10.5 | -8.8 | -10.7 | 6.3 | 5.8 | 4.6 | 4.3 | 4.4 | 1.4 | 3.8 | 4.8 | 4.8 | 4.6 | 3.3 | -24.5 | -10.7 | -12.6 | 1.1 | 4.7 | 3.4 | 3.0 |
| 1ZSH | -5.6 | -5.2 | -3.9 | -4.3 | -4.1 | -4.6 | 11.2 | 10.5 | 0.0 | -1.4 | 0.1 | -1.6 | 16.9 | 16.3 | 15.1 | 15.2 | 15.2 | 11.1 | 14.6 | 15.4 | 15.4 | 15.2 | 14.0 | -15.0 | -2.4 | -4.5 | 10.2 | 15.3 | 14.1 | 13.5 |
| 1G4M | -4.6 | -4.1 | -2.8 | -2.9 | -2.6 | -3.0 | 10.7 | 10.3 | 1.3 | 0.0 | 1.4 | -0.2 | 16.6 | 16.3 | 15.0 | 14.5 | 14.7 | 12.0 | 14.2 | 15.1 | 15.1 | 14.9 | 13.5 | -13.7 | -0.7 | -2.8 | 11.7 | 14.9 | 13.6 | 13.3 |
| 1JSY | -5.7 | -5.2 | -4.0 | -4.4 | -4.2 | -4.7 | 11.2 | 10.5 | -0.1 | -1.5 | 0.0 | -1.7 | 16.9 | 16.2 | 15.1 | 15.1 | 15.1 | 11.1 | 14.5 | 15.4 | 15.3 | 15.1 | 13.9 | -15.1 | -2.5 | -4.6 | 10.1 | 15.2 | 14.0 | 13.4 |
| 2WTR | -4.2 | -3.8 | -2.5 | -2.6 | -2.4 | -2.9 | 11.2 | 10.7 | 1.5 | 0.2 | 1.6 | 0.0 | 16.9 | 16.6 | 15.4 | 14.9 | 15.0 | 12.3 | 14.6 | 15.5 | 15.4 | 15.3 | 13.9 | -13.3 | -0.6 | -2.7 | 12.0 | 15.3 | 13.9 | 13.7 |
| 4JQI | -21.2 | -20.8 | -19.4 | -19.2 | -18.9 | -19.3 | -6.1 | -6.3 | -14.3 | -16.3 | -14.2 | -16.8 | 0.0 | -0.4 | -1.6 | -2.0 | -1.8 | -5.0 | -2.4 | -1.5 | -1.5 | -1.7 | -3.0 | -30.4 | -17.3 | -19.4 | -5.8 | -1.7 | -2.9 | -3.3 |
| 8J8Z | -21.1 | -20.6 | -19.5 | -20.2 | -20.1 | -20.7 | -5.3 | -5.8 | -16.2 | -17.2 | -16.1 | -17.3 | 0.4 | 0.0 | -1.2 | -1.6 | -1.5 | -4.5 | -2.0 | -1.1 | -1.1 | -1.3 | -2.6 | -30.5 | -18.8 | -21.0 | -5.0 | -1.2 | -2.5 | -2.9 |
| 8JAF | -20.0 | -19.6 | -18.3 | -18.3 | -18.0 | -18.3 | -4.1 | -4.6 | -13.8 | -15.4 | -13.7 | -15.6 | 1.6 | 1.1 | 0.0 | -0.3 | -0.2 | -3.5 | -0.8 | 0.1 | 0.1 | -0.1 | -1.4 | -29.0 | -16.0 | -18.0 | -4.2 | -0.1 | -1.3 | -1.7 |
| 6TKO | -19.6 | -19.2 | -18.0 | -18.9 | -18.8 | -19.5 | -3.7 | -4.2 | -14.8 | -15.8 | -14.7 | -16.0 | 2.0 | 1.6 | 0.4 | 0.0 | 0.2 | -3.1 | -0.4 | 0.5 | 0.5 | 0.3 | -1.0 | -29.0 | -17.6 | -19.9 | -3.7 | 0.4 | -0.9 | -1.3 |
| 6U1N | -19.5 | -19.1 | -17.9 | -18.7 | -18.6 | -19.3 | -3.9 | -4.4 | -14.4 | -15.6 | -14.3 | -15.9 | 1.8 | 1.4 | 0.3 | -0.2 | 0.0 | -3.3 | -0.6 | 0.4 | 0.4 | 0.2 | -1.1 | -29.1 | -17.5 | -19.9 | -4.0 | 0.2 | -1.0 | -1.4 |
| 6UP7 | -16.6 | -16.2 | -14.8 | -14.6 | -14.2 | -14.7 | -1.4 | -1.5 | -9.8 | -11.7 | -9.7 | -12.1 | 4.9 | 4.4 | 3.1 | 3.0 | 3.1 | 0.0 | 2.5 | 3.5 | 3.4 | 3.3 | 1.9 | -25.9 | -12.5 | -14.6 | -0.6 | 3.3 | 2.1 | 1.6 |
| 6PWC | -19.3 | -18.8 | -17.7 | -18.5 | -18.4 | -19.1 | -3.2 | -3.8 | -14.5 | -15.5 | -14.4 | -15.7 | 2.4 | 2.0 | 0.8 | 0.4 | 0.6 | -2.7 | 0.0 | 0.9 | 0.9 | 0.7 | -0.6 | -28.6 | -17.3 | -19.5 | -3.3 | 0.8 | -0.5 | -0.9 |
| 7R0C | -20.3 | -19.9 | -18.5 | -18.2 | -17.9 | -18.2 | -4.6 | -4.9 | -13.9 | -15.6 | -13.8 | -15.8 | 1.4 | 1.0 | -0.3 | -0.5 | -0.4 | -3.4 | -0.9 | 0.0 | 0.0 | -0.2 | -1.5 | -29.1 | -15.7 | -17.7 | -3.8 | -0.2 | -1.4 | -1.9 |
| 7SRS | -19.8 | -19.4 | -18.0 | -17.6 | -17.3 | -17.6 | -4.6 | -4.8 | -12.8 | -14.8 | -12.7 | -15.2 | 1.5 | 1.1 | -0.1 | -0.5 | -0.4 | -3.4 | -0.9 | 0.0 | 0.0 | -0.2 | -1.5 | -28.9 | -15.4 | -17.4 | -4.0 | -0.2 | -1.4 | -1.8 |
| 8WRZ | -19.9 | -19.5 | -18.1 | -17.7 | -17.4 | -17.7 | -4.4 | -4.7 | -13.3 | -15.0 | -13.2 | -15.3 | 1.7 | 1.3 | 0.0 | -0.3 | -0.2 | -3.2 | -0.7 | 0.2 | 0.2 | 0.0 | -1.3 | -29.1 | -15.2 | -17.1 | -3.6 | 0.0 | -1.2 | -1.6 |
| 8WU1 | -18.2 | -17.8 | -16.6 | -17.3 | -17.2 | -17.8 | -2.8 | -3.3 | -12.8 | -14.2 | -12.7 | -14.5 | 3.0 | 2.6 | 1.4 | 1.0 | 1.1 | -2.1 | 0.6 | 1.5 | 1.5 | 1.3 | 0.0 | -27.8 | -16.1 | -18.4 | -3.0 | 1.3 | 0.1 | -0.3 |
| 6K3F | 9.3 | 9.7 | 11.0 | 10.9 | 11.1 | 10.6 | 24.5 | 24.6 | 15.2 | 13.7 | 15.2 | 13.3 | 30.9 | 30.3 | 28.9 | 28.9 | 29.1 | 26.2 | 28.4 | 29.4 | 29.5 | 29.5 | 28.0 | 0.0 | 12.3 | 10.3 | 25.4 | 29.2 | 28.0 | 27.5 |
| 8J9K | -3.6 | -3.2 | -1.9 | -2.0 | -1.8 | -2.2 | 12.7 | 12.4 | 2.1 | 0.8 | 2.2 | 0.7 | 19.1 | 18.3 | 17.0 | 17.3 | 17.4 | 13.8 | 16.8 | 17.8 | 17.6 | 17.4 | 16.2 | -12.4 | 0.0 | -2.1 | 13.1 | 17.4 | 16.4 | 15.6 |
| 3P2D | -2.2 | -1.8 | -0.3 | 0.0 | 0.3 | -0.2 | 12.2 | 12.4 | 4.4 | 2.9 | 4.5 | 2.6 | 19.1 | 18.6 | 17.3 | 16.9 | 17.2 | 14.9 | 16.6 | 17.7 | 17.6 | 17.5 | 16.0 | -11.0 | 2.1 | 0.0 | 14.7 | 17.4 | 16.1 | 15.8 |
| 5TV1 | -15.7 | -15.3 | -14.1 | -14.7 | -14.5 | -15.0 | 0.7 | -0.2 | -10.6 | -11.7 | -10.5 | -11.8 | 5.9 | 5.5 | 4.4 | 3.9 | 4.1 | 0.6 | 3.5 | 4.4 | 4.3 | 4.1 | 2.9 | -25.1 | -12.8 | -14.8 | 0.0 | 4.3 | 2.9 | 2.6 |
| 8I10 | -19.5 | -19.1 | -17.9 | -18.3 | -18.1 | -18.7 | -4.3 | -4.7 | -13.6 | -15.2 | -13.5 | -15.7 | 1.7 | 1.2 | 0.1 | -0.4 | -0.2 | -3.6 | -0.8 | 0.2 | 0.2 | 0.0 | -1.3 | -28.9 | -17.0 | -19.2 | -4.4 | 0.0 | -1.2 | -1.6 |
| 8J8R | -18.5 | -18.1 | -16.7 | -16.2 | -15.9 | -16.2 | -3.2 | -3.5 | -11.5 | -13.4 | -11.4 | -13.8 | 2.9 | 2.5 | 1.3 | 0.9 | 1.0 | -2.0 | 0.5 | 1.4 | 1.4 | 1.2 | -0.1 | -27.4 | -13.8 | -15.8 | -2.5 | 1.2 | 0.0 | -0.4 |
| 8J8V | -18.0 | -17.6 | -16.2 | -15.9 | -15.6 | -15.9 | -2.7 | -3.0 | -11.2 | -13.1 | -11.1 | -13.5 | 3.3 | 2.9 | 1.7 | 1.3 | 1.4 | -1.7 | 0.9 | 1.8 | 1.8 | 1.6 | 0.3 | -27.1 | -13.6 | -15.6 | -2.3 | 1.6 | 0.4 | 0.0 |

**Table S4. Rotation angles among experimentally determined arrestin structures.**

The rotational angle refers to the rotation without projection onto the plane between the N and C domains. Note that for a given pair of structures, such as 4JQI and 1ZSH, the rotational angle is the same (23.1°) regardless of the order. Although these non-projected rotational angles generally have larger absolute values than the corresponding projected twist angles (Table S3), the differences in twist angles between certain βarr1 and βarr2 pairs are even greater than those observed in the corresponding rotational angles.

| | 3UGU | 4ZRG | 9C6E | 7JSM | 7JTB | 1CF1 | 4J2Q | 5W0P | 1ZSH | 1G4M | 1JSY | 2WTR | 4JQI | 8J8Z | 8JAF | 6TKO | 6U1N | 6UP7 | 6PWC | 7R0C | 7SRS | 8WRZ | 8WU1 | 6K3F | 8J9K | 3P2D | 5TV1 | 8I10 | 8J8R | 8J8V |
|---|---|---|---|---|---|---|---|---|---|---|---|---|---|---|---|---|---|---|---|---|---|---|---|---|---|---|---|---|---|---|
| 3UGU | 0.0 | 0.8 | 2.0 | 7.3 | 8.1 | 9.6 | 18.6 | 17.2 | 10.5 | 7.1 | 10.6 | 6.6 | 23.5 | 22.3 | 21.2 | 22.2 | 22.1 | 18.8 | 21.2 | 22.4 | 22.1 | 22.0 | 20.9 | 10.4 | 12.3 | 13.1 | 19.5 | 21.8 | 21.3 | 19.9 |
| 4ZRG | 0.8 | 0.0 | 2.0 | 7.8 | 8.7 | 10.1 | 17.8 | 16.5 | 10.9 | 7.3 | 11.0 | 6.7 | 22.9 | 21.7 | 20.6 | 21.5 | 21.4 | 18.4 | 20.6 | 21.8 | 21.5 | 21.4 | 20.3 | 10.9 | 12.7 | 13.6 | 19.1 | 21.2 | 20.6 | 19.3 |
| 9C6E | 2.0 | 2.0 | 0.0 | 6.2 | 7.0 | 8.5 | 17.6 | 16.0 | 9.0 | 5.3 | 9.1 | 4.8 | 22.1 | 20.8 | 19.8 | 20.9 | 20.7 | 17.2 | 19.9 | 20.9 | 20.7 | 20.5 | 19.6 | 11.7 | 10.9 | 12.1 | 17.7 | 20.4 | 19.9 | 18.5 |
| 7JSM | 7.3 | 7.8 | 6.2 | 0.0 | 1.0 | 2.3 | 21.9 | 19.5 | 5.9 | 3.5 | 6.0 | 3.7 | 24.4 | 23.2 | 22.5 | 23.6 | 23.3 | 18.1 | 22.5 | 23.2 | 23.0 | 22.7 | 22.3 | 11.9 | 5.0 | 6.0 | 17.2 | 23.0 | 22.5 | 21.2 |
| 7JTB | 8.1 | 8.7 | 7.0 | 1.0 | 0.0 | 1.7 | 22.5 | 20.0 | 5.3 | 3.7 | 5.4 | 4.2 | 24.7 | 23.5 | 22.8 | 24.0 | 23.7 | 18.4 | 22.9 | 23.5 | 23.4 | 23.1 | 22.7 | 12.5 | 4.4 | 5.4 | 17.3 | 23.4 | 23.0 | 21.6 |
| 1CF1 | 9.6 | 10.1 | 8.5 | 2.3 | 1.7 | 0.0 | 23.8 | 21.2 | 6.2 | 5.2 | 6.3 | 5.5 | 25.7 | 24.5 | 23.9 | 25.1 | 24.7 | 19.1 | 24.0 | 24.5 | 24.3 | 24.1 | 23.8 | 12.6 | 3.3 | 3.8 | 17.7 | 24.4 | 24.0 | 22.7 |
| 4J2Q | 18.6 | 17.8 | 17.6 | 21.9 | 22.5 | 23.8 | 0.0 | 4.1 | 21.2 | 19.0 | 21.2 | 18.3 | 9.1 | 8.1 | 6.8 | 7.4 | 7.8 | 12.0 | 7.0 | 8.9 | 8.5 | 8.7 | 6.9 | 27.9 | 24.3 | 26.4 | 15.9 | 7.4 | 7.6 | 6.5 |
| 5W0P | 17.2 | 16.5 | 16.0 | 19.5 | 20.0 | 21.2 | 4.1 | 0.0 | 19.0 | 16.7 | 19.0 | 15.8 | 7.1 | 6.2 | 4.8 | 5.3 | 5.4 | 8.1 | 4.5 | 6.3 | 5.9 | 5.9 | 4.3 | 26.3 | 21.4 | 23.4 | 11.9 | 5.2 | 5.0 | 3.6 |
| 1ZSH | 10.5 | 10.9 | 9.0 | 5.9 | 5.3 | 6.2 | 21.2 | 19.0 | 0.0 | 4.1 | 0.1 | 6.1 | 23.1 | 21.6 | 21.0 | 22.8 | 22.4 | 17.4 | 21.6 | 22.3 | 22.0 | 21.8 | 21.6 | 17.1 | 6.6 | 8.7 | 16.6 | 21.9 | 21.9 | 20.1 |
| 1G4M | 7.1 | 7.3 | 5.3 | 3.5 | 3.7 | 5.2 | 19.0 | 16.7 | 4.1 | 0.0 | 4.2 | 2.2 | 21.6 | 20.3 | 19.5 | 20.9 | 20.6 | 15.8 | 19.8 | 20.6 | 20.3 | 20.1 | 19.6 | 14.3 | 6.6 | 8.5 | 15.3 | 20.2 | 19.9 | 18.3 |
| 1JSY | 10.6 | 11.0 | 9.1 | 6.0 | 5.4 | 6.3 | 21.2 | 19.0 | 0.1 | 4.2 | 0.0 | 6.2 | 23.2 | 21.6 | 21.0 | 22.9 | 22.4 | 17.4 | 21.7 | 22.3 | 22.0 | 21.8 | 21.6 | 17.2 | 6.7 | 8.8 | 16.6 | 21.9 | 21.9 | 20.1 |
| 2WTR | 6.6 | 6.7 | 4.8 | 3.7 | 4.2 | 5.5 | 18.3 | 15.8 | 6.1 | 2.2 | 6.2 | 0.0 | 20.7 | 19.6 | 18.7 | 20.0 | 19.6 | 14.7 | 19.0 | 19.7 | 19.4 | 19.1 | 18.6 | 13.5 | 6.9 | 8.6 | 14.3 | 19.3 | 18.9 | 17.5 |
| 4JQI | 23.5 | 22.9 | 22.1 | 24.4 | 24.7 | 25.7 | 9.1 | 7.1 | 23.1 | 21.6 | 23.2 | 20.7 | 0.0 | 2.2 | 3.1 | 2.4 | 2.0 | 7.7 | 2.5 | 1.7 | 1.5 | 1.8 | 3.1 | 32.0 | 25.0 | 27.2 | 11.2 | 1.9 | 3.0 | 3.7 |
| 8J8Z | 22.3 | 21.7 | 20.8 | 23.2 | 23.5 | 24.5 | 8.1 | 6.2 | 21.6 | 20.3 | 21.6 | 19.6 | 2.2 | 0.0 | 1.4 | 3.4 | 2.9 | 7.5 | 2.7 | 3.0 | 2.4 | 2.7 | 3.4 | 31.2 | 24.0 | 26.2 | 11.1 | 2.0 | 3.7 | 2.9 |
| 8JAF | 21.2 | 20.6 | 19.8 | 22.5 | 22.8 | 23.9 | 6.8 | 4.8 | 21.0 | 19.5 | 21.0 | 18.7 | 3.1 | 1.4 | 0.0 | 3.1 | 2.8 | 7.5 | 2.2 | 3.3 | 2.7 | 2.9 | 2.8 | 30.0 | 23.4 | 25.7 | 11.3 | 1.7 | 3.4 | 1.9 |
| 6TKO | 22.2 | 21.5 | 20.9 | 23.6 | 24.0 | 25.1 | 7.4 | 5.3 | 22.8 | 20.9 | 22.9 | 20.0 | 2.4 | 3.4 | 3.1 | 0.0 | 0.7 | 7.8 | 1.3 | 1.9 | 1.9 | 2.1 | 1.3 | 30.5 | 24.7 | 26.7 | 11.6 | 1.5 | 1.3 | 2.8 |
| 6U1N | 22.1 | 21.4 | 20.7 | 23.3 | 23.7 | 24.7 | 7.8 | 5.4 | 22.4 | 20.6 | 22.4 | 19.6 | 2.0 | 2.9 | 2.8 | 0.7 | 0.0 | 7.2 | 1.0 | 1.2 | 1.1 | 1.4 | 1.2 | 30.5 | 24.2 | 26.3 | 11.0 | 1.1 | 1.1 | 2.5 |
| 6UP7 | 18.8 | 18.4 | 17.2 | 18.1 | 18.4 | 19.1 | 12.0 | 8.1 | 17.4 | 15.8 | 17.4 | 14.7 | 7.7 | 7.5 | 7.5 | 7.8 | 7.2 | 0.0 | 7.0 | 6.4 | 6.4 | 6.1 | 6.8 | 26.6 | 18.0 | 20.1 | 4.0 | 7.2 | 6.6 | 6.3 |
| 6PWC | 21.2 | 20.6 | 19.9 | 22.5 | 22.9 | 24.0 | 7.0 | 4.5 | 21.6 | 19.8 | 21.7 | 19.0 | 2.5 | 2.7 | 2.2 | 1.3 | 1.0 | 7.0 | 0.0 | 1.9 | 1.5 | 1.6 | 0.7 | 29.7 | 23.6 | 25.7 | 10.9 | 0.8 | 1.2 | 1.6 |
| 7R0C | 22.4 | 21.8 | 20.9 | 23.2 | 23.5 | 24.5 | 8.9 | 6.3 | 22.3 | 20.6 | 22.3 | 19.7 | 1.7 | 3.0 | 3.3 | 1.9 | 1.2 | 6.4 | 1.9 | 0.0 | 0.6 | 0.6 | 2.1 | 30.5 | 23.9 | 26.0 | 10.2 | 1.8 | 1.7 | 3.0 |
| 7SRS | 22.1 | 21.5 | 20.7 | 23.0 | 23.4 | 24.3 | 8.5 | 5.9 | 22.0 | 20.3 | 22.0 | 19.4 | 1.5 | 2.4 | 2.7 | 1.9 | 1.1 | 6.4 | 1.5 | 0.6 | 0.0 | 0.4 | 1.9 | 30.6 | 23.7 | 25.8 | 10.2 | 1.3 | 1.7 | 2.6 |
| 8WRZ | 22.0 | 21.4 | 20.5 | 22.7 | 23.1 | 24.1 | 8.7 | 5.9 | 21.8 | 20.1 | 21.8 | 19.1 | 1.8 | 2.7 | 2.9 | 2.1 | 1.4 | 6.1 | 1.6 | 0.6 | 0.4 | 0.0 | 2.0 | 30.5 | 23.4 | 25.5 | 9.8 | 1.5 | 1.7 | 2.5 |
| 8WU1 | 20.9 | 20.3 | 19.6 | 22.3 | 22.7 | 23.8 | 6.9 | 4.3 | 21.6 | 19.6 | 21.6 | 18.6 | 3.1 | 3.4 | 2.8 | 1.3 | 1.2 | 6.8 | 0.7 | 2.1 | 1.9 | 2.0 | 0.0 | 29.4 | 23.4 | 25.5 | 10.8 | 1.5 | 0.8 | 1.7 |
| 6K3F | 10.4 | 10.9 | 11.7 | 11.9 | 12.5 | 12.6 | 27.9 | 26.3 | 17.1 | 14.3 | 17.2 | 13.5 | 32.0 | 31.2 | 30.0 | 30.5 | 30.5 | 26.6 | 29.7 | 30.5 | 30.6 | 30.5 | 29.4 | 0.0 | 15.2 | 14.4 | 26.1 | 30.4 | 29.4 | 28.6 |
| 8J9K | 12.3 | 12.7 | 10.9 | 5.0 | 4.4 | 3.3 | 24.3 | 21.4 | 6.6 | 6.6 | 6.7 | 6.9 | 25.0 | 24.0 | 23.4 | 24.7 | 24.2 | 18.0 | 23.6 | 23.9 | 23.7 | 23.4 | 23.4 | 15.2 | 0.0 | 2.5 | 16.1 | 24.0 | 23.5 | 22.3 |
| 3P2D | 13.1 | 13.6 | 12.1 | 6.0 | 5.4 | 3.8 | 26.4 | 23.4 | 8.7 | 8.5 | 8.8 | 8.6 | 27.2 | 26.2 | 25.7 | 26.7 | 26.3 | 20.1 | 25.7 | 26.0 | 25.8 | 25.5 | 25.5 | 14.4 | 2.5 | 0.0 | 17.9 | 26.1 | 25.5 | 24.4 |
| 5TV1 | 19.5 | 19.1 | 17.7 | 17.2 | 17.3 | 17.7 | 15.9 | 11.9 | 16.6 | 15.3 | 16.6 | 14.3 | 11.2 | 11.1 | 11.3 | 11.6 | 11.0 | 4.0 | 10.9 | 10.2 | 10.2 | 9.8 | 10.8 | 26.1 | 16.1 | 17.9 | 0.0 | 11.0 | 10.5 | 10.2 |
| 8I10 | 21.8 | 21.2 | 20.4 | 23.0 | 23.4 | 24.4 | 7.4 | 5.2 | 21.9 | 20.2 | 21.9 | 19.3 | 1.9 | 2.0 | 1.7 | 1.5 | 1.1 | 7.2 | 0.8 | 1.8 | 1.3 | 1.5 | 1.5 | 30.4 | 24.0 | 26.1 | 11.0 | 0.0 | 1.9 | 1.9 |
| 8J8R | 21.3 | 20.6 | 19.9 | 22.5 | 23.0 | 24.0 | 7.6 | 5.0 | 21.9 | 19.9 | 21.9 | 18.9 | 3.0 | 3.7 | 3.4 | 1.3 | 1.1 | 6.6 | 1.2 | 1.7 | 1.7 | 1.7 | 0.8 | 29.4 | 23.5 | 25.5 | 10.5 | 1.9 | 0.0 | 2.3 |
| 8J8V | 19.9 | 19.3 | 18.5 | 21.2 | 21.6 | 22.7 | 6.5 | 3.6 | 20.1 | 18.3 | 20.1 | 17.5 | 3.7 | 2.9 | 1.9 | 2.8 | 2.5 | 6.3 | 1.6 | 3.0 | 2.6 | 2.5 | 1.7 | 28.6 | 22.3 | 24.4 | 10.2 | 1.9 | 2.3 | 0.0 |

**Table S5. Subsegment definitions for the PIA analysis.**

The subsegments were defined by integrating those previously described (Hirsch et al. 1999, Gurevich et al. 2018), with minor adjustments based on visual inspection of experimentally determined structures, and by subdividing long β-strands into shorter subsegments to improve resolution (ensuring that each subsegment in an extended β-strand conformation contains no more than nine residues).

| id | arr1 | | arr2 | | arr3 | | name |
|---|---|---|---|---|---|---|---|
| | start | end | start | end | start | end | |
| **b1** | 12 | 16 | 8 | 12 | 9 | 13 | β-strand I |
| **b2** | 23 | 26 | 19 | 22 | 20 | 23 | β-strand II |
| **b3** | 30 | 33 | 26 | 29 | 27 | 30 | β-strand III |
| **b4** | 41 | 46 | 37 | 42 | 38 | 43 | β-strand IV |
| **b5i** | 56 | 62 | 52 | 58 | 53 | 59 | β-strand V-i |
| **b5ii** | 63 | 68 | 59 | 64 | 60 | 65 | β-strand V-ii |
| **fl** | 69 | 78 | 65 | 74 | 66 | 75 | finger loop |
| **b6i** | 79 | 84 | 75 | 80 | 76 | 81 | β-strand VI-i |
| **b6ii** | 85 | 89 | 81 | 85 | 82 | 86 | β-strand VI-ii |
| **as1** | 90 | 102 | 86 | 99 | 87 | 100 | arrestin switch I (ASwI) |
| **3e** | 103 | 112 | 100 | 109 | 101 | 110 | 3-element helix |
| **b7** | 115 | 120 | 112 | 117 | 113 | 118 | β-strand VII |
| **b8** | 130 | 132 | 127 | 129 | 128 | 130 | β-strand VIII |
| **ml** | 133 | 141 | 130 | 138 | 131 | 139 | middle/139 loop |
| **b9i** | 142 | 148 | 139 | 145 | 140 | 146 | β-strand IX-i |
| **b9ii** | 149 | 154 | 146 | 151 | 147 | 152 | β-strand IX-ii |
| **157l** | 155 | 169 | 152 | 163 | 153 | 164 | 157 loop |
| **b10** | 170 | 178 | 164 | 172 | 165 | 173 | β-strand X |
| **hin** | 179 | 190 | 173 | 182 | 174 | 183 | hinge/ASwII |
| **b11** | 191 | 197 | 183 | 189 | 184 | 190 | β-strand XI/ASwII |
| **b12** | 202 | 209 | 196 | 203 | 197 | 204 | β-strand XII |
| **b13** | 213 | 215 | 207 | 209 | 208 | 210 | β-strand XIII |
| **b14** | 220 | 228 | 214 | 222 | 215 | 223 | β-strand XIV |
| **b15i** | 234 | 240 | 228 | 234 | 229 | 235 | β-strand XV-i |
| **b15ii** | 241 | 247 | 235 | 241 | 236 | 242 | β-strand XV-ii |
| **cl** | 248 | 252 | 242 | 246 | 243 | 247 | C loop |
| **b16i** | 253 | 258 | 247 | 252 | 248 | 253 | β-strand XVI-i |
| **b16ii** | 259 | 264 | 253 | 258 | 254 | 259 | β-strand XVI-ii |
| **b17** | 272 | 280 | 266 | 274 | 267 | 275 | β-strand XVII |
| **ll0** | 284 | 295 | 278 | 289 | 279 | 290 | lariat loop-0 |
| **ll1** | 296 | 303 | 290 | 297 | 291 | 298 | lariat loop-1 |
| **as3** | 313 | 322 | 307 | 316 | 308 | 317 | ASwIII |
| **b18i** | 323 | 329 | 317 | 323 | 318 | 324 | β-strand XVIII-i |
| **b18ii** | 330 | 335 | 324 | 329 | 325 | 330 | β-strand XVIII-ii |
| **344l** | 336 | 346 | 330 | 342 | 331 | 335 | 344 loop |
| **b19i** | 347 | 351 | 343 | 347 | 336 | 340 | β-strand IXX-i |
| **b19ii** | 352 | 356 | 348 | 352 | 341 | 345 | β-strand IXX-ii |

### SUPPLEMENTAL FIGURES

**Figure S1. The scopes of temperatures visited in the 100-replica (a) and 200-replica (b) TREMD simulations.**

The simulations were carried out with 100 and 200 replicas using 100 and 200 6-GPU nodes. The analysis was conducted on the unsorted simulation results. For each 80 ns run, the temperature scope explored (i.e., the difference between the highest and lowest visited temperatures) is shown for the first 40 ns (orange) and the full 80 ns (blue), plotted against the starting temperature of each unsorted replica trajectory.

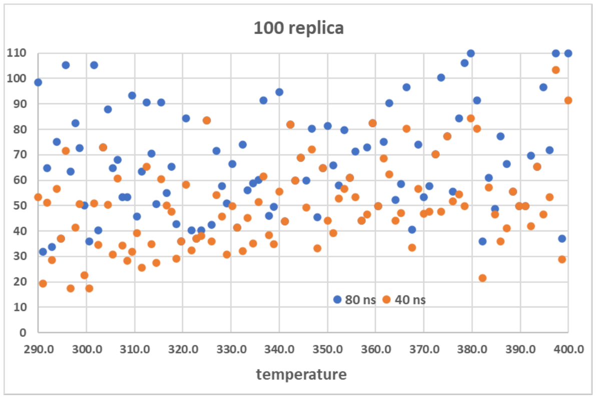


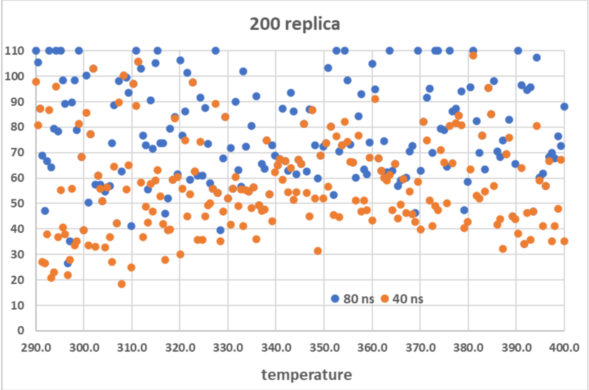

**Figure S2. The partial dissociation of V2Rpp observed in the 200-replica TREMD run, but not in either classical MD simulations or the 100-replica TREMD run.**

The results shown are from the 300.5K resorted trajectory. After 70 ns, one frame shows the dissociation - the dotted arrows indicate P1, P2, and P3, the first three phosphorylated residues in V2Rpp, are dissociated from the extended groove. Note that P4 did not make strong interaction with the N domain. Thus, the entire P1-P4 region was dissociated. This portion of V2Rpp was reassociated back to the groove in the following simulations.

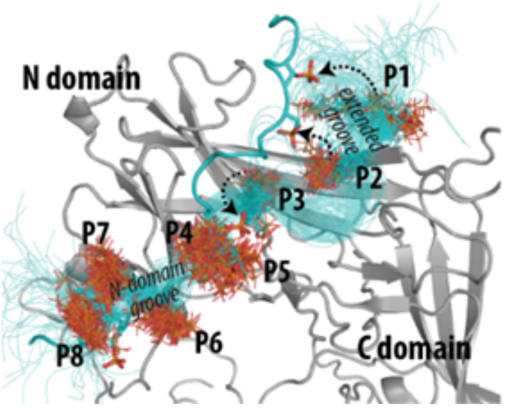

**Figure S3. Comparison of three superposition schemes in calculating RMSDs of MD simulations.**

The simulated conditions with or without the application of HMR are indicated on the right. For each condition, both structures 1ZSH (basal, left) and 4JQI (active, right) are used for three schemes, (1) aligning the entire main body to calculate its RMSD (termed the "main-body" scheme), (2) aligning the N domain to calculate the C-domain RMSD ("N=>C"), and (3) aligning the C domain to calculate the N-domain RMSD ("C=>N"). The peak values of each distribution are labeled and indicated with red dotted lines.

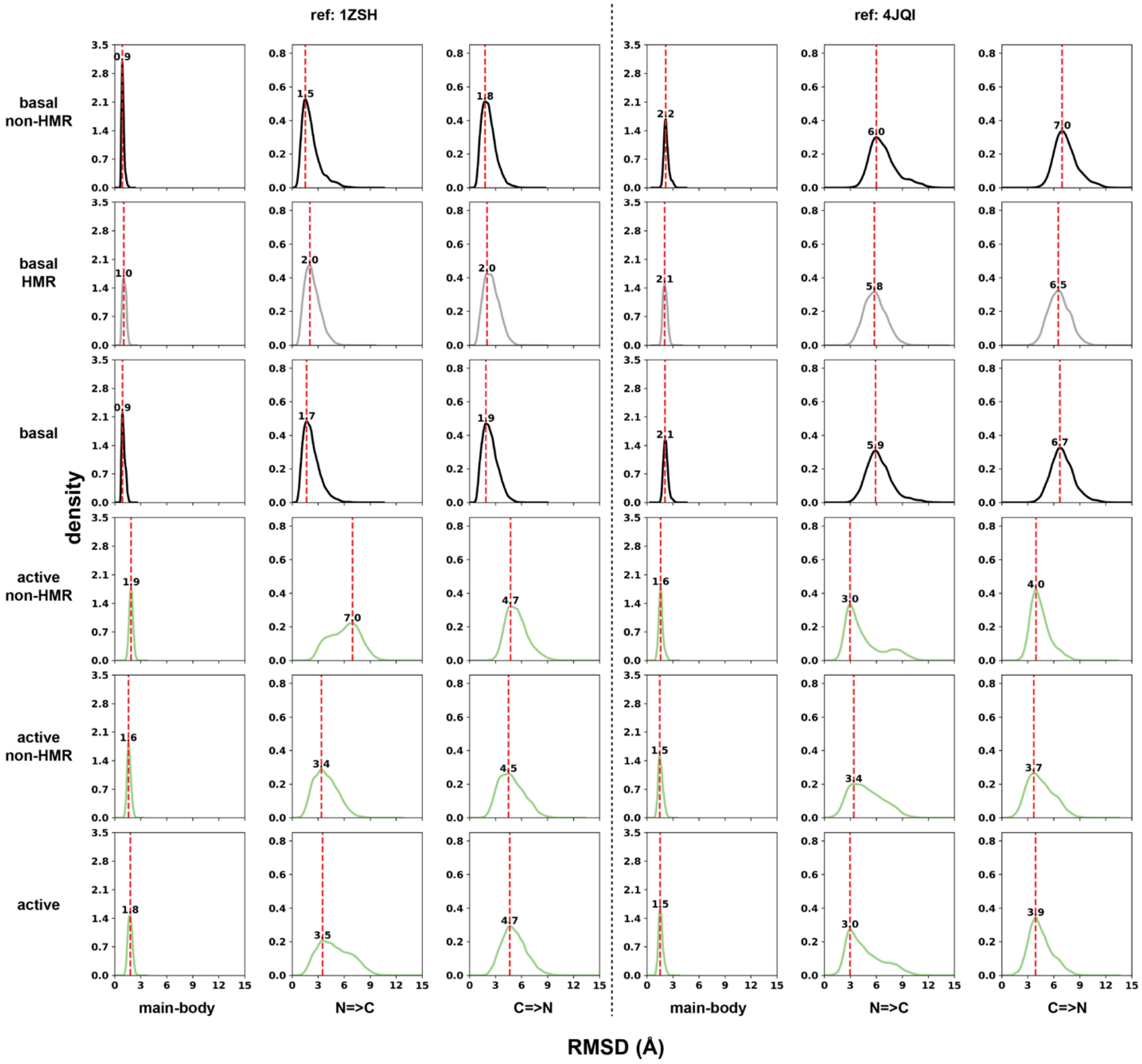

**Figure S4. Comparison of three superposition schemes in calculating RMSDs of TREMD simulations.**

The simulated conditions with or without the application of HMR are indicated on the right. For each condition, both structures 1ZSH (basal, left) and 4JQI (active, right) are used for three schemes, (1) aligning the entire main body to calculate its RMSD (termed the “main-body” scheme), (2) aligning the N domain to calculate the C-domain RMSD (“N=>C”), and (3) aligning the C domain to calculate the N-domain RMSD (“C=>N”). The peak values of each distribution are labeled and indicated with red dotted lines.

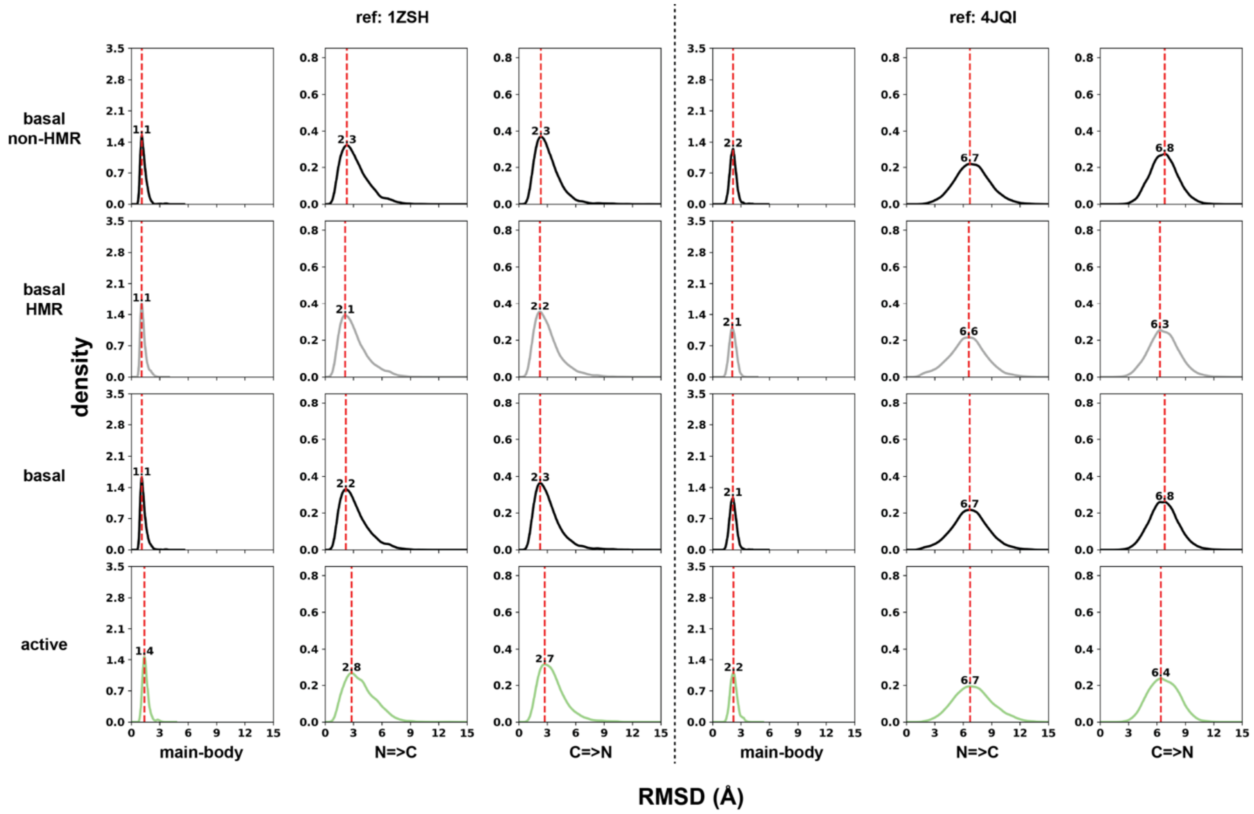

**Figure S5. The definition of the twist angle.**

Panels A and B illustrate the N and C domains as schematic spheres, with arrows connecting their centers of mass (COMs). The NC plane is defined as the plane perpendicular to the axis extending from the COM of the N domain to that of the C domain in the reference structure, which is the “basal” state in this scheme. (Note that if the arrow instead points from the N-domain COM to the C-domain COM of the active structure, then the active structure serves as the reference.) From panel A to panel B, the viewing angle rotates from a side view to a view along the N-to-C axis. The rotational angle ($\gamma$) is defined in rotational space as the angle extracted from the quaternion representation of the rotation matrix that best maps the selected Cα atoms of the C domain in the target structure onto those of the reference, following alignment of their N domains (see Methods). The twist angle is the projection of this rotational angle onto the NC plane. Panel C shares the same viewing direction as panel B and shows the selected Cα atoms in the N domain (gray) and in the C domain (green for the basal state, magenta for the active state), which are used to calculate the COMs. The larger, darker-colored spheres mark the resulting COMs.

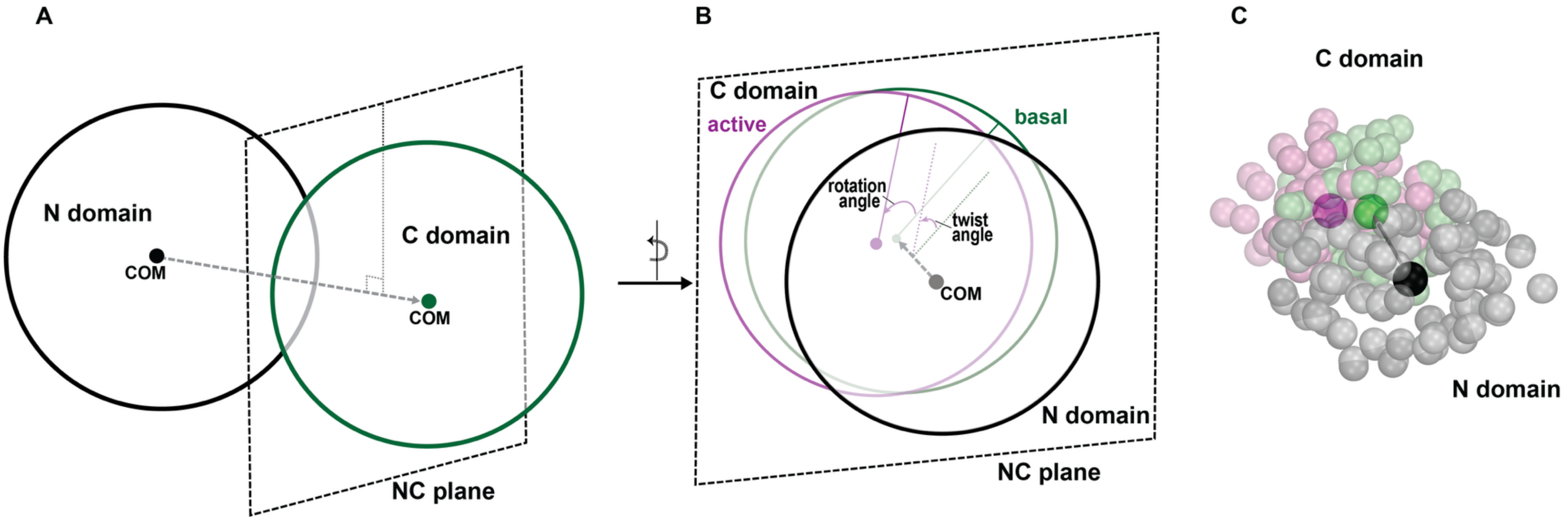
A
N domain
COM
C domain
COM
NC plane
B
C domain
active
basal
rotation angle
twist angle
COM
N domain
NC plane
C
C domain
N domain

**Figure S6. Multiple sequence alignment of arrestins highlighting conserved and divergent regions.**

The alignment includes sequences from three species (bovine, human, and rat) for each of three arrestin isoforms: arr1 (visual arrestin or ARRS), arr2 (β-arrestin-1 or ARRB1), and arr3 (β-arrestin-2 or ARRB2). While the N and C domains are highly conserved across isoforms and species, notable sequence divergence is observed in some of the functionally important regions, which are boxed. These include the 157 loop, implicated in phosphopeptide engagement, and the C-tail, which varies in both length and composition among isoforms. The C-tail is subdivided into proximal, middle, and distal segments, as indicated by differently shaded green bars. Notably, the extra motif is absent in arr1, and sequence divergence is also apparent at the junction between the extra and chameleon motifs when comparing arr2 and arr3.

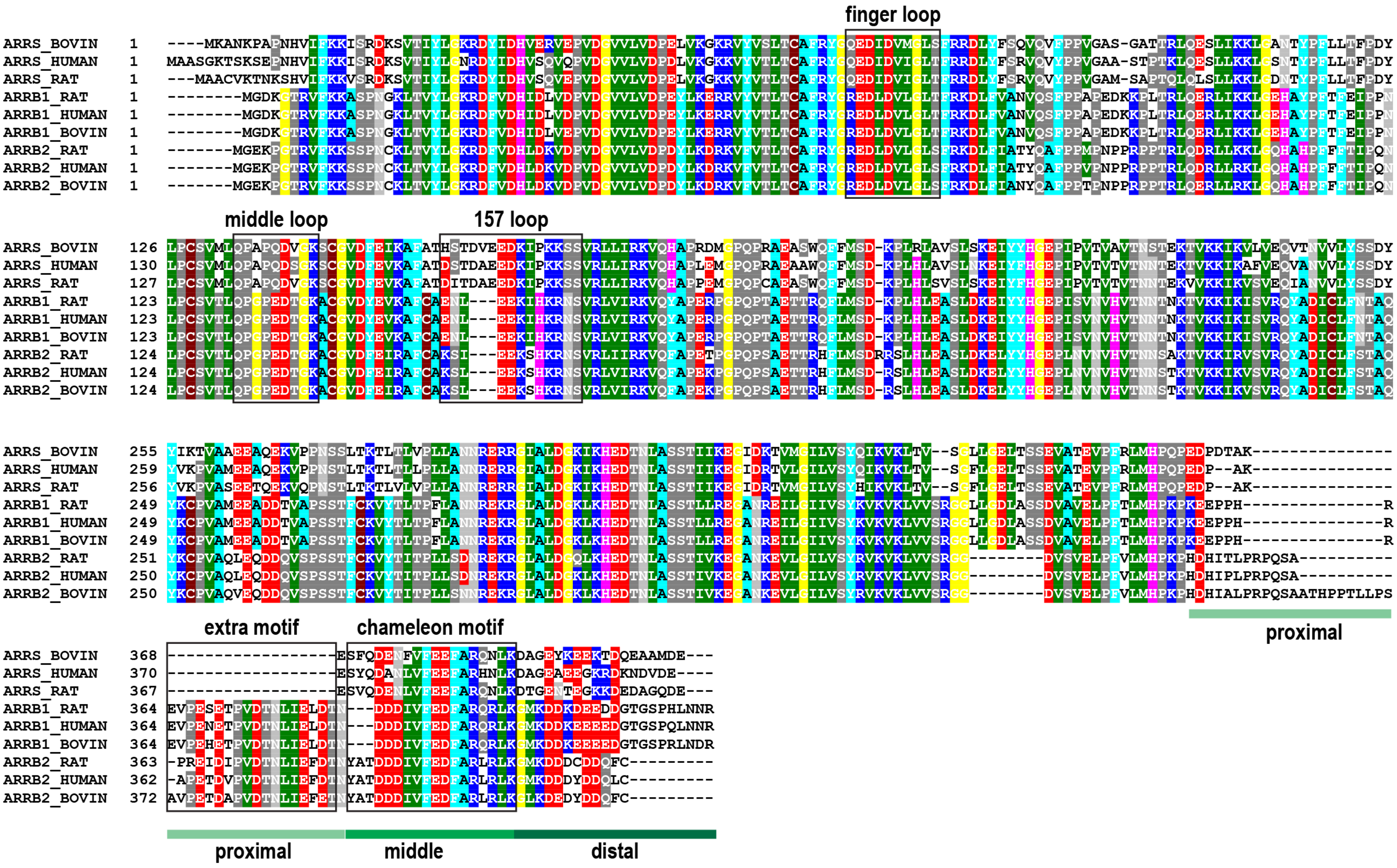
finger loop
middle loop
157 loop
proximal
extra motif
chameleon motif
proximal
middle
distal
ARRS_BOVIN
ARRS_HUMAN
ARRS_RAT
ARRB1_RAT
ARRB1_HUMAN
ARRB1_BOVIN
ARRB2_RAT
ARRB2_HUMAN
ARRB2_BOVIN

**Figure S7. The interactions of the βarr1 tail with the main body in the V2Rpp-bound state.**

The heatmap shows the normalized contact ratios between residues 357-418 of the βarr1 tail and residues 1-356 of the main body, calculated over all frames of the 312.9K replica in the V2Rpp-bound TREMD simulation. Each pixel represents the contact ratio of a tail-main body residue pair, with more blueish colors indicating higher contact frequency. Local interaction hotspots, identified by applying a 5×5 maximum filter and a contact ratio threshold of 0.1, are marked with red dots. These hotspots were used to define anchor residue pairs for downstream clustering of tail conformations (see Fig. 6). Among residues C-terminal to position 366 of the βarr1 tail, two distinct interaction patches were identified: residues 370-376 and 392-396, which frequently contact ASwIII (residues 307-316) and the C-loop (residues 242-246) of the C domain, respectively. Notably, the 370-376 patch also frequently interacted with the residues in the loop between β-strands VII and VIII (residues 118-126) of the N domain.

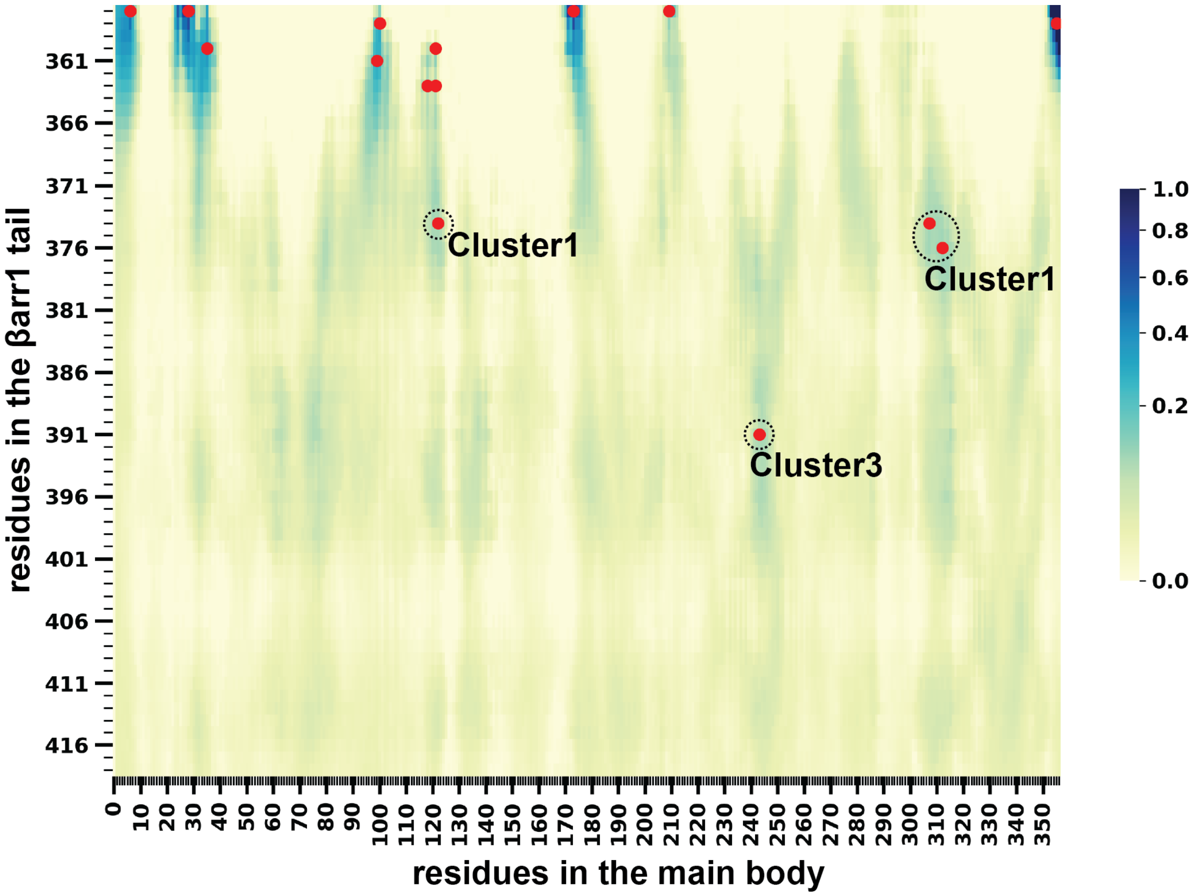

### REFERENCE

Asher, W. B., D. S. Terry, G. G. A. Gregorio, A. W. Kahsai, A. Borgia, B. Xie, A. Modak, Y. Zhu, W. Jang, A. Govindaraju, L. Y. Huang, A. Inoue, N. A. Lambert, V. V. Gurevich, L. Shi, R. J. Lefkowitz, S. C. Blanchard and J. A. Javitch (2022). "GPCR-mediated beta-arrestin activation deconvoluted with single-molecule precision." Cell **185**(10): 1661-1675 e1616.

Bous, J., A. Fouillen, H. Orcel, S. Trapani, X. Cong, S. Fontanel, J. Saint-Paul, J. Lai-Kee-Him, S. Urbach, N. Sibille, R. Sounier, S. Granier, B. Mouillac and P. Bron (2022). "Structure of the vasopressin hormone-V2 receptor-beta-arrestin1 ternary complex." Sci Adv **8**(35): eabo7761.

Cao, C., X. Barros-Alvarez, S. Zhang, K. Kim, M. A. Damgen, O. Panova, C. M. Suomivuori, J. F. Fay, X. Zhong, B. E. Krumm, R. H. Gumpper, A. B. Seven, M. J. Robertson, N. J. Krogan, R. Huttenhain, D. E. Nichols, R. O. Dror, G. Skiniotis and B. L. Roth (2022). "Signaling snapshots of a serotonin receptor activated by the prototypical psychedelic LSD." Neuron **110**(19): 3154-3167 e3157.

Chen, Q., N. A. Perry, S. A. Vishnivetskiy, S. Berndt, N. C. Gilbert, Y. Zhuo, P. K. Singh, J. Tholen, M. D. Ohi, E. V. Gurevich, C. A. Brautigam, C. S. Klug, V. V. Gurevich and T. M. Iverson (2017). "Structural basis of arrestin-3 activation and signaling." Nat Commun **8**(1): 1427.

Dobson, C. M., A. Sali and M. Karplus (1998). "Protein Folding: A Perspective from Theory and Experiment." Angew Chem Int Ed Engl **37**(7): 868-893.

Essmann, U., L. Perera, M. L. Berkowitz, T. Darden, H. Lee and L. G. Pedersen (1995). "A smooth particle mesh Ewald method." J Chem Phys **103**(19): 8577-8593.

Feller, S. E., Y. H. Zhang, R. W. Pastor and B. R. Brooks (1995). "Constant-Pressure Molecular-Dynamics Simulation - the Langevin Piston Method." Journal of Chemical Physics **103**(11): 4613-4621.

Granzin, J., A. Cousin, M. Weirauch, R. Schlesinger, G. Buldt and R. Batra-Safferling (2012). "Crystal structure of p44, a constitutively active splice variant of visual arrestin." J Mol Biol **416**(5): 611-618.

Granzin, J., A. Stadler, A. Cousin, R. Schlesinger and R. Batra-Safferling (2015). "Structural evidence for the role of polar core residue Arg175 in arrestin activation." Sci Rep **5**: 15808.

Gurevich, V. V., E. V. Gurevich and V. N. Uversky (2018). "Arrestins: structural disorder creates rich functionality." Protein Cell **9**(12): 986-1003.

Han, M., V. V. Gurevich, S. A. Vishnivetskiy, P. B. Sigler and C. Schubert (2001). "Crystal structure of beta-arrestin at 1.9 A: possible mechanism of receptor binding and membrane Translocation." Structure **9**(9): 869-880.

Hirsch, J. A., C. Schubert, V. V. Gurevich and P. B. Sigler (1999). "The 2.8 A crystal structure of visual arrestin: a model for arrestin's regulation." Cell **97**(2): 257-269.

Hopkins, C. W., S. Le Grand, R. C. Walker and A. E. Roitberg (2015). "Long-Time-Step Molecular Dynamics through Hydrogen Mass Repartitioning." J Chem Theory Comput **11**(4): 1864-1874.

Huang, J., S. Rauscher, G. Nawrocki, T. Ran, M. Feig, B. L. de Groot, H. Grubmuller and A. D. MacKerell, Jr. (2017). "CHARMM36m: an improved force field for folded and intrinsically disordered proteins." Nat Methods **14**(1): 71-73.

Huang, W., M. Masureel, Q. Qu, J. Janetzko, A. Inoue, H. E. Kato, M. J. Robertson, K. C. Nguyen, J. S. Glenn, G. Skiniotis and B. K. Kobilka (2020). "Structure of the neurotensin receptor 1 in complex with beta-arrestin 1." Nature **579**(7798): 303-308.

Jo, S., T. Kim, V. G. Iyer and W. Im (2008). "CHARMM-GUI: a web-based graphical user interface for CHARMM." J Comput Chem **29**(11): 1859-1865.

Jorgensen, W. L., J. Chandrasekhar, J. D. Madura, R. W. Impey and M. L. Klein (1983). "Comparison of Simple Potential Functions for Simulating Liquid Water." Journal of Chemical Physics **79**(2): 926-935.

Karplus, M. (2011). "Behind the folding funnel diagram." Nat Chem Biol **7**(7): 401-404.

Kim, Y. J., K. P. Hofmann, O. P. Ernst, P. Scheerer, H. W. Choe and M. E. Sommer (2013). "Crystal structure of pre-activated arrestin p44." Nature **497**(7447): 142-146.
Kofke, D. A. (2002). "On the acceptance probability of replica-exchange Monte Carlo trials." The Journal of Chemical Physics **117**(15): 6911-6914.
Latorraca, N. R., M. Masureel, S. A. Hollingsworth, F. M. Heydenreich, C. M. Suomivuori, C. Brinton, R. J. L. Townshend, M. Bouvier, B. K. Kobilka and R. O. Dror (2020). "How GPCR Phosphorylation Patterns Orchestrate Arrestin-Mediated Signaling." Cell.
Latorraca, N. R., J. K. Wang, B. Bauer, R. J. L. Townshend, S. A. Hollingsworth, J. E. Olivieri, H. E. Xu, M. E. Sommer and R. O. Dror (2018). "Molecular mechanism of GPCR-mediated arrestin activation." Nature **557**(7705): 452-456.
Lee, Y., T. Warne, R. Nehme, S. Pandey, H. Dwivedi-Agnihotri, M. Chaturvedi, P. C. Edwards, J. Garcia-Nafria, A. G. W. Leslie, A. K. Shukla and C. G. Tate (2020). "Molecular basis of beta-arrestin coupling to formoterol-bound beta1-adrenoceptor." Nature **583**(7818): 862-866.
Liao, Y. Y., H. Zhang, Q. Shen, C. Cai, Y. Ding, D. D. Shen, J. Guo, J. Qin, Y. Dong, Y. Zhang and X. M. Li (2023). "Snapshot of the cannabinoid receptor 1-arrestin complex unravels the biased signaling mechanism." Cell **186**(26): 5784-5797 e5717.
Maharana, J., F. K. Sano, P. Sarma, M. K. Yadav, L. Duan, T. M. Stepniewski, M. Chaturvedi, A. Ranjan, V. Singh, S. Saha, G. Mahajan, M. Chami, W. Shihoya, J. Selent, K. Y. Chung, R. Banerjee, O. Nureki and A. K. Shukla (2024). "Molecular insights into atypical modes of beta-arrestin interaction with seven transmembrane receptors." Science **383**(6678): 101-108.
Maharana, J., P. Sarma, M. K. Yadav, S. Saha, V. Singh, S. Saha, M. Chami, R. Banerjee and A. K. Shukla (2023). "Structural snapshots uncover a key phosphorylation motif in GPCRs driving beta-arrestin activation." Mol Cell **83**(12): 2091-2107 e2097.
Milano, S. K., Y. M. Kim, F. P. Stefano, J. L. Benovic and C. Brenner (2006). "Nonvisual arrestin oligomerization and cellular localization are regulated by inositol hexakisphosphate binding." J Biol Chem **281**(14): 9812-9823.
Milano, S. K., H. C. Pace, Y. M. Kim, C. Brenner and J. L. Benovic (2002). "Scaffolding functions of arrestin-2 revealed by crystal structure and mutagenesis." Biochemistry **41**(10): 3321-3328.
Min, K., H. J. Yoon, J. Y. Park, M. Baidya, H. Dwivedi-Agnihotri, J. Maharana, M. Chaturvedi, K. Y. Chung, A. K. Shukla and H. H. Lee (2020). "Crystal Structure of beta-Arrestin 2 in Complex with CXCR7 Phosphopeptide." Structure **28**(9): 1014-1023 e1014.
Neale, C., R. Pomes and A. E. Garcia (2016). "Peptide Bond Isomerization in High-Temperature Simulations." J Chem Theory Comput **12**(4): 1989-1999.
Neria, E., S. Fischer and M. Karplus (1996). "Simulation of activation free energies in molecular systems." Journal of Chemical Physics **105**(5): 1902-1921.
Ngo, V. A., Y. T. Lin and D. Perez (2023). "Improving Estimation of the Koopman Operator with Kolmogorov-Smirnov Indicator Functions." J Chem Theory Comput **19**(20): 7187-7198.
Noe, F. and S. Fischer (2008). "Transition networks for modeling the kinetics of conformational change in macromolecules." Curr Opin Struct Biol **18**(2): 154-162.
Periole, X. and A. E. Mark (2007). "Convergence and sampling efficiency in replica exchange simulations of peptide folding in explicit solvent." J Chem Phys **126**(1): 014903.
Phillips, J. C., R. Braun, W. Wang, J. Gumbart, E. Tajkhorshid, E. Villa, C. Chipot, R. D. Skeel, L. Kale and K. Schulten (2005). "Scalable molecular dynamics with NAMD." J Comput Chem **26**(16): 1781-1802.
Phillips, J. C., D. J. Hardy, J. D. C. Maia, J. E. Stone, J. V. Ribeiro, R. C. Bernardi, R. Buch, G. Fiorin, J. Henin, W. Jiang, R. McGreevy, M. C. R. Melo, B. K. Radak, R. D. Skeel, A. Singharoy, Y. Wang, B. Roux, A. Aksimentiev, Z. Luthey-Schulten, L. V. Kale, K. Schulten, C. Chipot and E. Tajkhorshid (2020). "Scalable molecular dynamics on CPU and GPU architectures with NAMD." J Chem Phys **153**(4): 044130.

Rosta, E. and G. Hummer (2009). "Error and efficiency of replica exchange molecular dynamics simulations." J Chem Phys **131**(16): 165102.
Salom, D., P. D. Kiser and K. Palczewski (2025). "Insights into the Activation and Self-Association of Arrestin-1." Biochemistry **64**(2): 364-376.
Sanbonmatsu, K. Y. and A. E. Garcia (2002). "Structure of Met-enkephalin in explicit aqueous solution using replica exchange molecular dynamics." Proteins **46**(2): 225-234.
Sander, C. L., J. Luu, K. Kim, D. Furkert, K. Jang, J. Reichenwallner, M. Kang, H. J. Lee, B. T. Eger, H. W. Choe, D. Fiedler, O. P. Ernst, Y. J. Kim, K. Palczewski and P. D. Kiser (2022). "Structural evidence for visual arrestin priming via complexation of phosphoinositols." Structure **30**(2): 263-277 e265.
Shukla, A. K., A. Manglik, A. C. Kruse, K. Xiao, R. I. Reis, W. C. Tseng, D. P. Staus, D. Hilger, S. Uysal, L. Y. Huang, M. Paduch, P. Tripathi-Shukla, A. Koide, S. Koide, W. I. Weis, A. A. Kossiakoff, B. K. Kobilka and R. J. Lefkowitz (2013). "Structure of active beta-arrestin-1 bound to a G-protein-coupled receptor phosphopeptide." Nature **497**(7447): 137-141.
Staus, D. P., H. Hu, M. J. Robertson, A. L. W. Kleinhenz, L. M. Wingler, W. D. Capel, N. R. Latorraca, R. J. Lefkowitz and G. Skiniotis (2020). "Structure of the M2 muscarinic receptor-beta-arrestin complex in a lipid nanodisc." Nature.
Stein, A. and T. Kortemme (2013). "Improvements to robotics-inspired conformational sampling in rosetta." PLoS One **8**(5): e63090.
Wang, Y., L. Wu, T. Wang, J. Liu, F. Li, L. Jiang, Z. Fan, Y. Yu, N. Chen, Q. Sun, Q. Tan, T. Hua and Z. J. Liu (2024). "Cryo-EM structure of cannabinoid receptor CB1-beta-arrestin complex." Protein Cell **15**(3): 230-234.
Webb, B. and A. Sali (2016). "Comparative Protein Structure Modeling Using MODELLER." Curr Protoc Bioinformatics **54**: 5 6 1-5 6 37.
Yin, W., Z. Li, M. Jin, Y. L. Yin, P. W. de Waal, K. Pal, Y. Yin, X. Gao, Y. He, J. Gao, X. Wang, Y. Zhang, H. Zhou, K. Melcher, Y. Jiang, Y. Cong, X. Edward Zhou, X. Yu and H. Eric Xu (2019). "A complex structure of arrestin-2 bound to a G protein-coupled receptor." Cell Res **29**(12): 971-983.
Yoo, J. and A. Aksimentiev (2012). "Improved Parametrization of Li+, Na+, K+, and Mg2+ Ions for All-Atom Molecular Dynamics Simulations of Nucleic Acid Systems." The Journal of Physical Chemistry Letters **3**(1): 45-50.
Zhan, X., L. E. Gimenez, V. V. Gurevich and B. W. Spiller (2011). "Crystal structure of arrestin-3 reveals the basis of the difference in receptor binding between two non-visual subtypes." J Mol Biol **406**(3): 467-478.
Zhang, W., C. Wu and Y. Duan (2005). "Convergence of replica exchange molecular dynamics." J Chem Phys **123**(15): 154105.
Zhou, X. E., X. Gao, A. Barty, Y. Kang, Y. He, W. Liu, A. Ishchenko, T. A. White, O. Yefanov, G. W. Han, Q. Xu, P. W. de Waal, K. M. Suino-Powell, S. Boutet, G. J. Williams, M. Wang, D. Li, M. Caffrey, H. N. Chapman, J. C. Spence, P. Fromme, U. Weierstall, R. C. Stevens, V. Cherezov, K. Melcher and H. E. Xu (2016). "X-ray laser diffraction for structure determination of the rhodopsin-arrestin complex." Sci Data **3**: 160021.
Zhou, X. E., Y. He, P. W. de Waal, X. Gao, Y. Kang, N. Van Eps, Y. Yin, K. Pal, D. Goswami, T. A. White, A. Barty, N. R. Latorraca, H. N. Chapman, W. L. Hubbell, R. O. Dror, R. C. Stevens, V. Cherezov, V. V. Gurevich, P. R. Griffin, O. P. Ernst, K. Melcher and H. E. Xu (2017). "Identification of Phosphorylation Codes for Arrestin Recruitment by G Protein-Coupled Receptors." Cell **170**(3): 457-469 e413.
